# Bacterial vampirism mediated through taxis to serum

**DOI:** 10.1101/2023.07.07.548164

**Authors:** Siena J. Glenn, Zealon Gentry-Lear, Michael Shavlik, Michael J. Harms, Thomas J. Asaki, Arden Baylink

## Abstract

Bacteria of the family Enterobacteriaceae are associated with gastrointestinal (GI) bleeding and bacteremia and are a leading cause of death, from sepsis, for individuals with inflammatory bowel diseases. The bacterial behaviors and mechanisms underlying why these bacteria are prone to bloodstream entry remains poorly understood. Herein, we report that clinical isolates of non-typhoidal *Salmonella enterica* serovars, *Escherichia coli*, and *Citrobacter koseri* are rapidly attracted toward sources of human serum. To simulate GI bleeding, we utilized a custom injection-based microfluidics device and found that femtoliter volumes of human serum are sufficient to induce the bacterial population to swim toward and aggregate at the serum source. This response is orchestrated through chemotaxis, and a major chemical cue driving chemoattraction is L-serine, an amino acid abundant in serum that is recognized through direct binding by the chemoreceptor Tsr. We report the first crystal structures of *Salmonella* Typhimurium Tsr in complex with L-serine and identify a conserved amino acid recognition motif for L-serine shared among Tsr orthologues. By mapping the phylogenetic distribution of this chemoreceptor we found Tsr to be widely conserved among Enterobacteriaceae and numerous World Health Organization priority pathogens associated with bloodstream infections. Lastly, we find that Enterobacteriaceae use human serum as a source of nutrients for growth and that chemotaxis and the chemoreceptor Tsr provides a competitive advantage for migration into enterohaemorrhagic lesions. We term this bacterial behavior of taxis toward serum, colonization of hemorrhagic lesions, and the consumption of serum nutrients, as “bacterial vampirism” which may relate to the proclivity of Enterobacteriaceae for bloodstream infections.

## Introduction

Bacteria use chemosensory systems to survey and navigate their environments ^1–3^. In the exceptionally dynamic environment of the host gut, peristalsis and flow constantly perturb the microscopic physicochemical landscape. Responding to such transient and shifting stimuli is enabled by chemosensing that allows bacteria to rapidly restructure their populations, within seconds, through taxis toward or away from effector sources ^4–6^. Enteric pathogens and pathobionts use chemosensing to colonize specific tissues, based on sources of exogenous nutrients, toxins, and host-emitted cues, through control of motility and adhesion systems such as chemotaxis, chemokinesis, twitching, and biofilm formation ^3,7–11^. Previously, we determined that most bacterial genera classified by the World Health Organization (WHO) as ‘priority pathogens’ employ sophisticated chemosensory systems for efficient infection and exert precise control over their colonization topography ^3,12,13^. These include multidrug-resistant Enterobacteriaceae pathogens of the gut such as *Salmonella enterica*, Enterohemorrhagic *Escherichia coli* (EHEC), *Citrobacter koseri*, and *Enterobacter cloacae*, which present major challenges for nosocomial infections and global health ^3,13^.

Mapping the chemosensing signals enteric bacteria perceive within the host can provide insight into pathogenesis and perhaps be a source of drug targets for inhibiting pathogen navigation and colonization ^3,14^. Gut dysbiosis—a diseased host state induced by pathologies, inflammation and infections—exposes enteric bacteria to chemical stimuli distinct from that of a healthy gut that could encourage opportunistic pathogenesis ^3,15^. Yet, the stimuli sensed by bacterial pathogens and pathobionts within the dysbiotic gut environ environment remain poorly understood. Investigating how chemosensing controls the responses of pathogen populations to chemical features associated with dysbiosis presents an opportunity to gain deeper insights into the critical juncture between infection resolution and exacerbation (Fig. 1) ^3,15–17^.

**Fig. 1.**
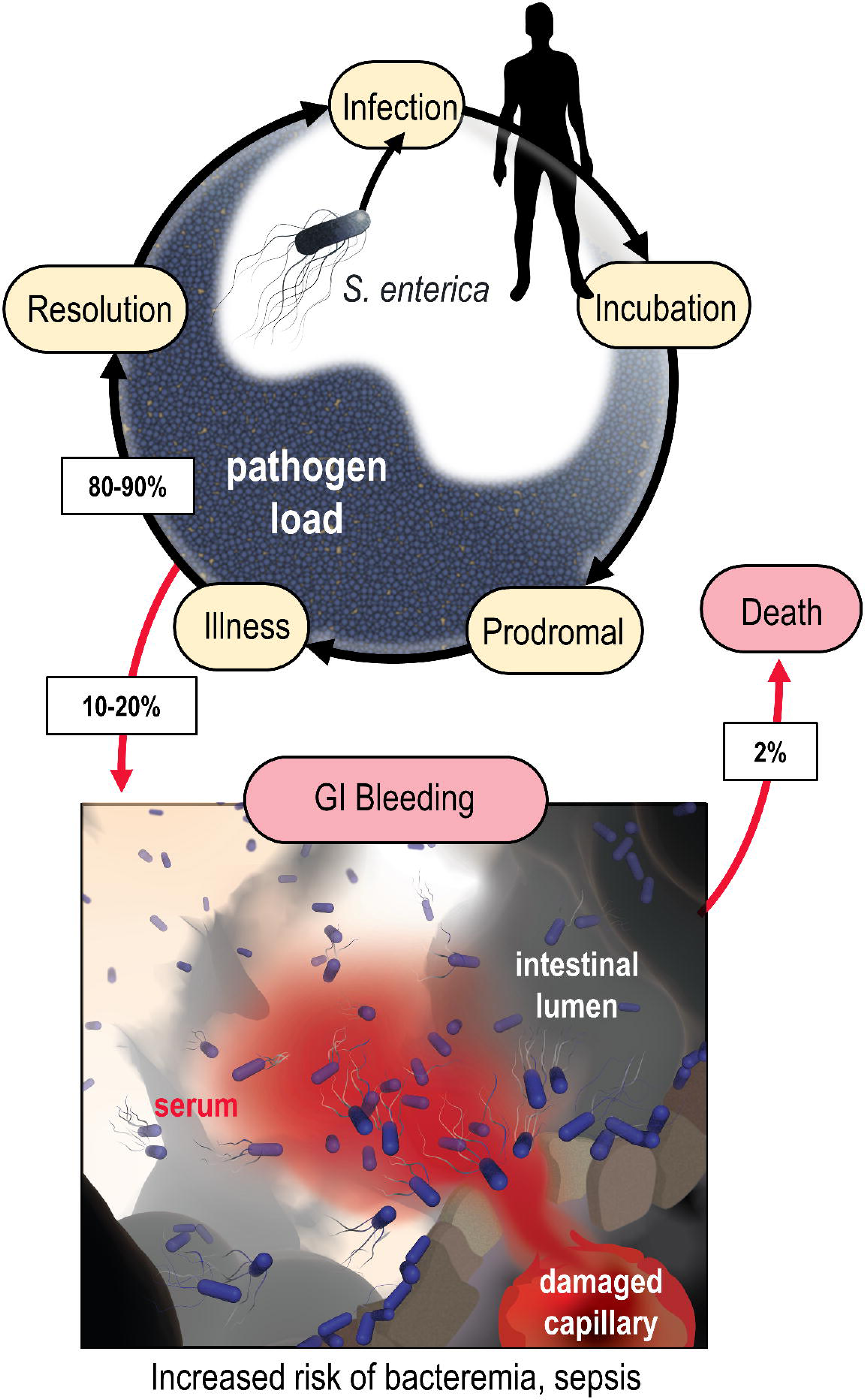
Model of the microenvironment of bacterial-induced hemorrhagic lesions. The typical course of non-typhoidal *S. enterica* infections is shown proceeding through infection, incubation, prodromal, illness, and resolution stages (black arrows). The atypical route of GI bleeding, associated with increased mortality and morbidity, is shown in red arrows, with rates approximated from available literature. An artistic depiction of bacterial injury tropism is shown bottom.

In a diseased gut, enteric bacteria may encounter a distinctive host-derived chemical feature not found in a healthy gut: GI bleeding. The microenvironment of an enteric hemorrhagic lesion involves a source of serum, the liquid component of blood, emanating from the host tissue and diffusing into the intestinal lumen (Fig. 1). In effect, this creates a microscopic gradient, i.e., a microgradient, of chemicals flowing outward from a point source that may serve as chemosensory signals for bacteria. Enteric infections caused by Enterobacteriaceae species can lead to GI bleeding, a condition with a significant risk of mortality. Although severe GI bleeding isn’t a typical outcome of enteric infection, it is a substantial burden on human health, affecting around 40-150 out of every 100,000 individuals annually, with a fatality rate ranging from 6% to 30% (Fig. 1) ^3,13,18–20,20–23^. Enterobacteriaceae are prone to bloodstream entry and are a leading cause of sepsis-related deaths in individuals with inflammatory bowel diseases (IBD) that have recurrent enterohemorrhagic lesions ^24,25^. Intestinal intra-abdominal abscesses, microperforations, and fistulas associated with IBD predispose patients to GI bleeding and bacteremia ^26,27^. Despite the established connection between Enterobacteriaceae-induced sepsis and GI bleeding, it remained unknown whether these bacteria perceive serum through chemosensing.

Serum is a complex biological solution with components that may enhance, or hinder, bacterial growth. It offers a rich reservoir of nutrients for bacteria: sugars and amino acids at millimolar concentrations and essential metals like iron and zinc ^3,28^. Yet, serum also contains host factors that inhibit bacterial proliferation in the bloodstream such as cathelicidin, defensins, and the complement system ^29,30^. Consequently, how enteric pathogens and pathobionts might respond to serum diffusing into a liquid environment remained unclear. To address this open question, we elucidated how Enterobacteriaceae species use chemosensing to respond to serum, the molecular mechanism of this response, and how chemosensing is employed to enter and migrate into enterohemorrhagic lesions. Across all examined scenarios we observe these bacteria to exhibit remarkable sensitivity and attraction towards human serum. These findings suggest that environmental stimuli unique to the dysbiotic gut, sensed through bacterial chemosensory systems, can encourage pathogenic behaviors and adverse consequences for the host.

## Results

### Use of the chemosensory injection rig assay (CIRA) to study polymicrobial chemosensing behaviors

To model features of enteric bleeding *in vitro*, we utilized an experimental system to inject minute quantities of human serum into a pond of motile bacteria and observe real-time responses by microscopy (Fig. 2A). The system and methodology, which we refer to herein as the chemosensory injection rig assay (CIRA), offers several advantages for studying bacterial chemosensing and localization in response to serum: (1) we can use bona fide human serum, (2) the readouts are direct measurements of real-time localization dynamics of the bacterial population, and (3) similar to a bleeding event, a source of fresh serum is continuously emitted. By employing multichannel fluorescence imaging of differentially labeled bacterial populations we observe polymicrobial interactions through head-to-head comparisons of bacterial behavior within the same experiment.

**Fig. 2.**
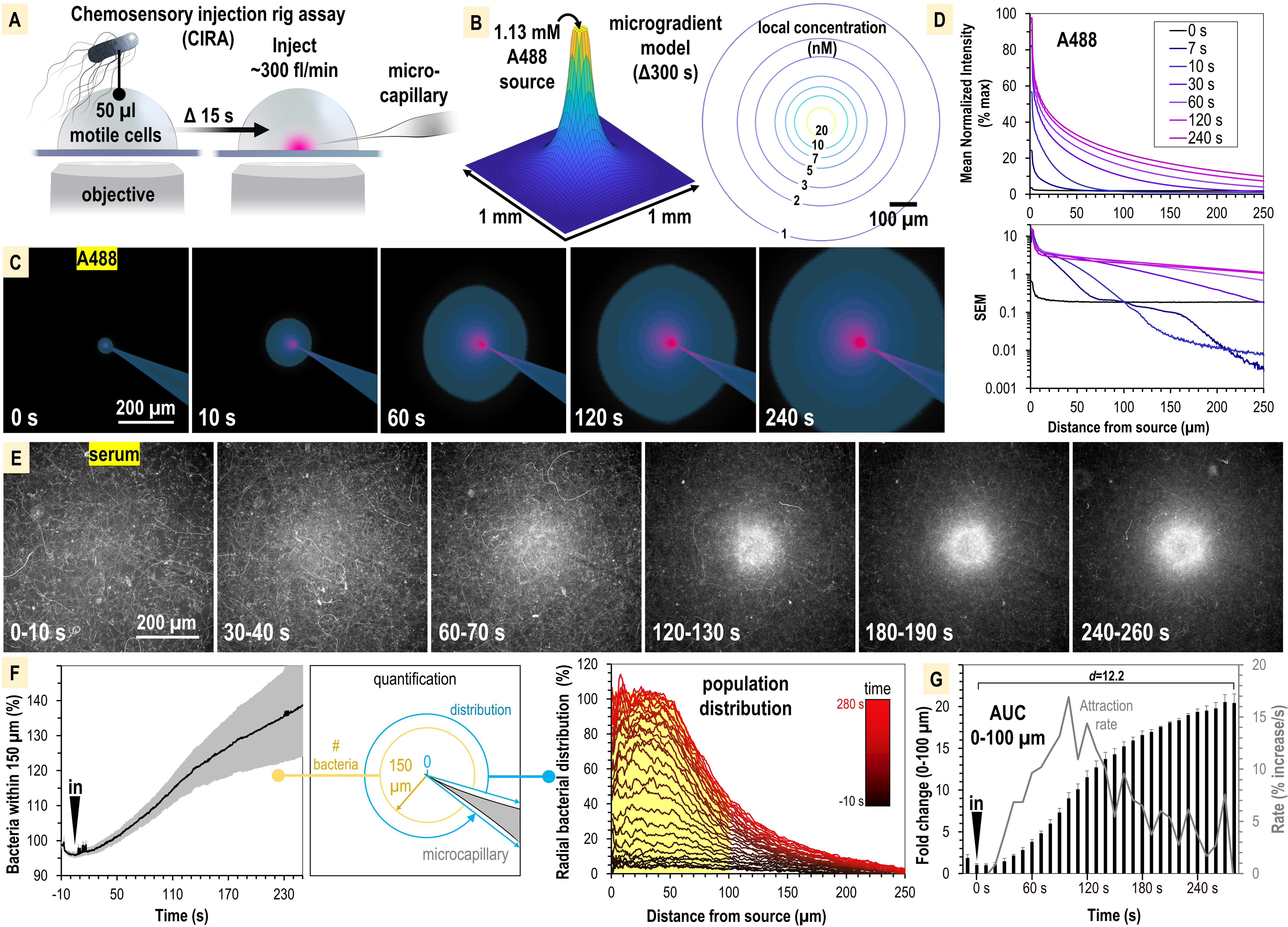
*S. enterica* Typhimurium IR715 rapidly localizes toward human serum. A. CIRA experimental design. B. CIRA microgradient diffusion model, simulated with a source of 1.13 mM A488 dye after 300 s of injection. C. Experimental visualization of the CIRA microgradient with A488 dye. D. Injection and diffusion of A488 dye. Shown top is the mean normalized fluorescence intensity at representative time points as a function of distance from the source, and shown bottom is the standard error of the mean (SEM) for these data (n=6). E. Response of *S. enterica* Typhimurium IR715 to human serum (max projections over 10 s intervals). F. Quantification of *S. enterica* Typhimurium IR715 attraction response to human serum (n=4, 37° C) characterized as either the relative number of bacteria within 150 µm of the source (left), or the radial distribution of the bacterial population over time (right, shown in 10 s intervals). G. Area under the curve (AUC) versus time for the bacterial population within 100 µm of the serum treatment source (area indicated in yellow in panel F). Effect size (Cohen’s *d*) between the treatment start and endpoints is indicated. Insertion of the treatment microcapillary is indicated with black ‘in’ arrow. Attraction rate over time indicated in gray. Data shown are means, error bars indicate SEM.

The CIRA microgradient is established through injection at a constant rate of ∼300 femtoliters per minute, through a 0.5 µm glass microcapillary, and expands over time through diffusion (Fig. 2A-B, Fig. S1). The diffusion of serum metabolites can be reasonably approximated by the three-dimensional differential diffusion equation (Fig. 2B, Fig. S1, Fig. S2, Method Details). We calculate that the introduction of a minute volume of novel effector produces a steep microgradient, in which a millimolar source recedes to nanomolar concentrations after diffusion across only a few hundred microns (Fig. 2B, Fig. S1). For instance, we expect a bacterium 100 µm from a 1 mM infinite-volume source to experience a local concentration of a small molecule effector (diffusion coefficient approximately 4 x 10^−6^ cm^2^ s^-1^) of 10 nM after 300 s of injection (Fig. 2B, Fig. S1). To visualize the microgradient experimentally, we utilized Alexa Fluor 488 dye (A488) and observed the microgradient to be stable and consistent across replicates (Fig. 2C-D, Movie 1). As a test case of the accuracy of the microgradient modeling, we compared our calculated values for A488 diffusion to the normalized fluorescence intensity at time 120 s. We determined the concentration to be accurate within 5% over the distance range 70-270 µm (Fig. S2). At smaller distances (<70 µm) the measured concentration is approximately 10% lower than that predicted by the computation. This could be due to advection effects near the injection site that would tend to enhance the effective local diffusion rate. Additionally, we found no behavioral differences in treatments with buffer in the range of pH 4-9, indicating small, localized pH changes are inconsequential for taxis in our system, and that any artifactual forces, such as flow, account for only minor changes to bacterial distribution at the population level, in the range of ±10% (Fig. S1).

### Non-typhoidal S. enterica serovars exhibit rapid attraction to human serum

We first studied the chemosensing behaviors of *S.* Typhimurium IR715, which is a derivative of ATCC 14028 originally isolated from domesticated chickens, and is used extensively in the study of *Salmonella* pathogenesis (Table S1)^31–34^. As a treatment, we utilized commercially available human serum that was not heat-inactivated nor exposed to chemical or physical treatments that would be expected to alter its native complement properties (see Materials & Methods). We assessed the response of motile IR715, containing a fluorescent mPlum marker, to a source of human serum over the course of 5 minutes (Fig. 2E, Movie 1). During this timeframe, we witnessed a rapid attraction response whereby the motile bacterial population reorganized from a random distribution to one concentrated within a 100-150 µm radius of the serum source (Fig. 2E-F, Movie 1). To compare responses between CIRA experiments, we plotted data as either the relative number of bacteria within 150 µm of the treatment (Fig. 2F, left panel) or a radial distribution of the population (Fig. 2F, right panel) over time. By these metrics, we determined *S.* Typhimurium IR715 is attracted to human serum. The bacterial population in proximity to the serum source doubles by 40 s, reaches a maximal attraction rate by 90 s, and approaches equilibrium by 300 s post-treatment (Fig. 2G).

To test whether serum attraction is observed in *Salmonella* strains that infect humans, and if responses differ among non-typhoidal *Salmonella* serovars, we employed dual-channel CIRA imaging to compete *S.* Typhimurium IR715 against representative clinical isolates of Typhimurium (SARA1), Newport (M11018046001A), and Enteriditis (05E01375). These serovars are the most common in North American infections ^35^. In varying magnitude, all strains showed attraction responses to human serum, seen as a significant distribution bias toward the serum source relative to the experiment periphery (Fig. 3, Movie 2). We noted that in some experiments the population peak is 50-75 µm from the source, possibly due to a compromise between achieving proximity to nutrients in the serum and avoidance of bactericidal serum elements, but this behavior was not consistent across all experiments. Overall, our data show *S. enterica* serovars that cause disease in humans are exquisitely sensitive to human serum, responding to femtoliter quantities as an attractant, and that distinct reorganization at the population level occurs within minutes of exposure (Fig. 3, Movie 2).

**Fig. 3.**
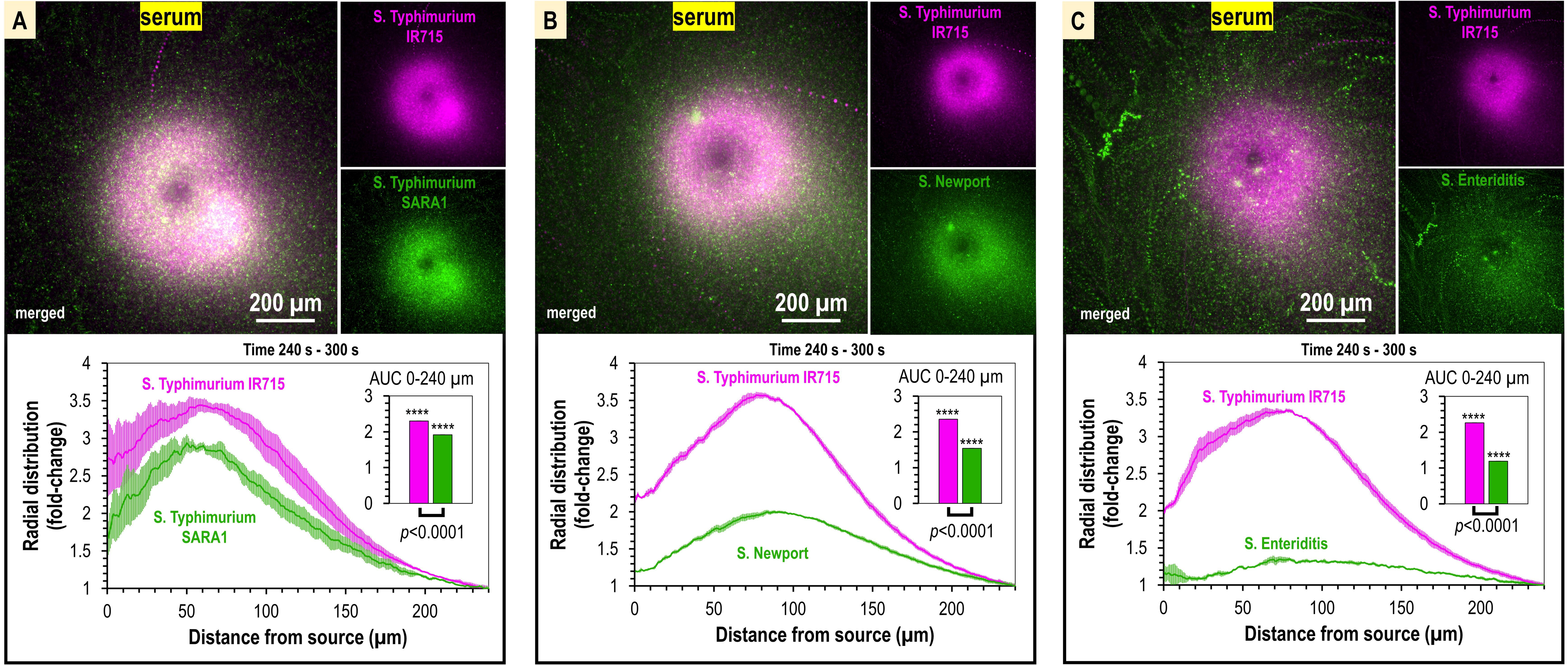
Taxis to human serum is retained across *S. enterica* clinical isolates and diverse serovars. A-C. CIRA competition experiments between *S.* Typhimurium IR715 (pink) and clinical isolates (green) responding to human serum for 5 mins (n=4, 37° C). Images are representative max projections over the final minute of treatment. Radial distributions calculated from max projections and averaged across replicates are shown as fold-change relative to the image periphery at 240 µm from the source. Inset plots show fold-change AUC of strains in the same experiment relative to an expected baseline of 1 (no change). *p*-values shown are calculated with an unpaired two-sided t-test comparing the means of the two strains, or one-sided t-test to assess statistical significance in terms of change from 1-fold (stars). Data shown are means, error bars indicate SEM.

### Chemotaxis and the chemoreceptor Tsr mediate serum attraction

Serum is a complex biological solution that contains sugars, amino acids, and other metabolites that could serve as attractant signals ^3^. Based on the rapid attraction of motile, swimming bacteria to the treatment source, characteristic of chemotactic behaviors ^5^, we hypothesized that one or more of these chemical components are specifically recognized as chemoattractants through the repertoire of chemoreceptors possessed by *Salmonellae* (Fig. 4A). Based on known chemoreceptor-chemoattractant ligand interactions ^3,36^, we identified three chemoreceptors that might mediate taxis toward serum: (1) taxis to serine and repellents (Tsr), which responds to L- serine, and reportedly also norepinephrine (NE) and 3,4-dihydroxymandelic acid (DHMA); (2) taxis to ribose and glucose/galactose (Trg), which responds to glucose and galactose; and (3) taxis to aspartate and repellents (Tar), which responds to L-aspartate (Fig. 4A). We modeled the local concentration profile of these effectors based on their typical concentrations in human serum (Fig. 4B). Of these, by far the two most prevalent chemoattractants in serum are glucose (5 mM) and L-serine (100-300 µM) (Fig. 4B-F). This suggested to us that the chemoreceptors Trg and/or Tsr could play important roles in serum attraction. These chemoreceptors were also previously shown to provide colonization advantages during *S.* Typhimurium infection ^34^.

**Fig. 4.**
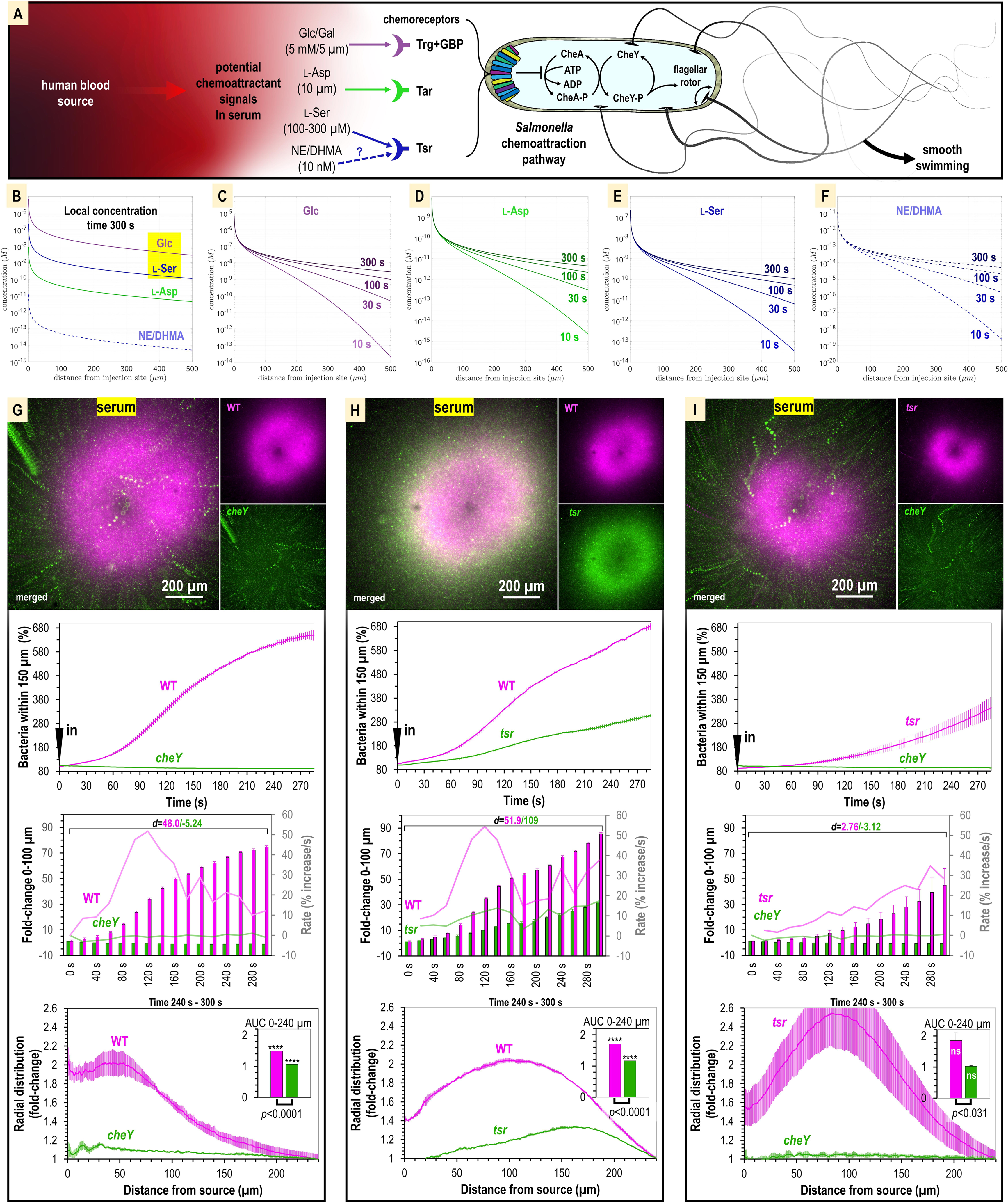
Attraction to human serum is mediated through chemotaxis and the chemoreceptor Tsr. A. Potential mechanisms involved in *Salmonella* sensing of chemoattractants present in human serum. Approximate concentrations of these effectors in human serum are indicated in parentheses ^3^. B-F. Microgradient modeling of serum chemoattractant concentrations. G-I. CIRA competition experiments between *S.* Typhimurium IR715 WT and isogenic mutants *cheY*, or *tsr*, and *tsr* versus *cheY,* in response to human serum (n=3-4, 37° C). Rates in terms of fold-change are indicated with light pink/light green lines and plotted on the gray secondary y-axis. Inset AUC plots are shown as described in Fig. 3. Data are means and error bars are SEM.

To test the role of chemotaxis in serum attraction, we competed wildtype (WT) *S.* Typhimurium IR715 against a chemotaxis-null isogenic *cheY* mutant, which possesses swimming motility but is blind to chemoeffector signals (Fig. 4G, Movie 3). Whereas the WT mounts a robust attraction response to serum, the *cheY* mutant population remains randomly distributed (Fig. 4G, Movie 3). We also observed the *cheY* mutant to exhibit a slight decline in cells proximal to the treatment source over time, which we attribute to cellular crowding effects from the influx of WT cells (Fig. 4G, Movie 3). In this background, the fraction of WT cells within 100 µm of the serum source increases by 70-fold, with the maximal rate of attraction achieved by 120 s post-treatment. Thus, we determined that chemotaxis is namely responsible for the rapid localization of *S. enterica* to human serum.

We next analyzed strains with deletions of the chemoreceptor genes *trg,* or *tsr,* to test the roles of these chemoreceptors in mediating taxis to serum. We were surprised to find that the *trg* strain had deficiencies in swimming motility (data not shown). This was not noted in earlier work but could explain the severe infection disadvantage of this mutant ^34^. Because motility is a prerequisite for chemotaxis, we chose not to study the *trg* mutant further, and instead focused our investigations on Tsr. We compared chemoattraction responses in dual-channel CIRA experiments between the WT and *tsr* mutant and observed an interesting behavior whereby both strains exhibited chemoattraction, but the *tsr* mutant distribution was relegated to a halo at the periphery of the WT peak (Fig. 4H, Movie 3). The WT efficiently outcompetes the *tsr* mutant such that by 5 minutes post-treatment the ratio of WT to *tsr* cells proximal to the serum source is 3:1 (Fig. 4H, Movie 3). Similar to *cheY*, we presume the *tsr* halo results from cellular crowding effects induced by the high density of WT cells near the serum source. To test how the *tsr* mutant responds to serum in the absence of a strong competitor, we compared chemoattraction between *tsr* and *cheY*. In this background, *tsr* chemoattraction remained diminished relative to that of WT, but the *tsr* distribution shifted closer to the serum source (Fig. 4I, Movie 3). Since *tsr* mutation diminishes serum attraction but does not eliminate it, we conclude that multiple chemoattractant signals and chemoreceptors mediate taxis to serum. To further understand the mechanism of this behavior we chose to focus on Tsr as a representative chemoreceptor involved in the response, presuming that serum taxis involves one, or more, of the chemoattractants recognized by Tsr that is present in serum: L-serine, NE, or DHMA.

### S. Typhimurium exhibits chemoattraction to L-serine, but not NE or DHMA

We next sought to identify the specific chemoattractants driving Tsr-mediated serum attraction. Suspecting that one effector might be L-serine, we took advantage of the fact that Tsr can only bind the L and not the D enantiomer and treated the serum with serine racemase, an enzyme that converts L- to D-serine. The attraction of *S.* Typhimurium IR715 to serum treated with serine racemase is diminished and the population is more diffuse (Fig. S3A-B). In competition experiments between WT and chemotaxis-deficient strains, the WT cells proximal to the treatment source are reduced by about half (Fig. S3C), and WT no longer possesses an attraction advantage over the *tsr* mutant (Fig. S3D). These data support that the deficiency of the *tsr* strain in serum taxis is due to an inability to sense L-serine within serum.

We next used CIRA to examine responses to purified effectors diluted in buffer. The concentration of L-serine in human serum ranges from approximately 100-400 µm, depending on diet and health factors ^37–39^, whereas the neurotransmitter NE, and its metabolized form, DHMA, are thought to circulate at approximately 10 nM ^40,41^. It has been proposed that, like L-serine, NE and/or DHMA are sensed directly by Tsr and at even higher (nanomolar) affinity ^42,43^. We used CIRA to test the response of *S.* Typhimurium IR715 to L-serine, NE, and DHMA. We observed robust chemoattraction responses to L-serine, evident by the accumulation of cells toward the treatment source (Fig. S3E, Movie 4), but no response to NE or DHMA, with the cells remaining randomly distributed even after 5 minutes of exposure (Fig. S3F-I, Movie 5, Movie S1). Following extensive investigation and adjusting experiment parameters such as culture protocol, cell density, temperature, injection flow, and compound concentration, we saw no evidence that NE or DHMA are chemoeffectors for *S.* Typhimurium (Fig. S3F-I, Movie 5, Movie S1). Since this result is surprising given previous reports ^42,43^, we have provided a database of 17 videos of CIRA experiments showing the null response to DHMA and NE, versus L-serine, under various conditions (Data S1). Together, our data indicate the chemoattractant component of serum that is sensed through Tsr is L-serine.

Chemoattraction to L-serine has mostly been studied in the context of model laboratory strains and has not been rigorously evaluated for *S. enterica* clinical isolates or various serovars. To establish whether L-serine sensing is observed in strains responsible for human infections, we used dual-channel CIRA to compare chemoattraction to L-serine between *S.* Typhimurium IR715 and clinical isolates (Fig. 5A-C, Movie 4). In each case we observe robust chemoattraction, though there are differences in sensitivity to L-serine. The magnitude of chemoattraction was highest for *S.* Typhimurium SARA1 and *S.* Newport, whereas *S.* Typhimurium IR715 and *S.* Enteriditis showed lower responses, which could relate to the different host specificities of these serovars and strains (Fig. 5A-C, Movie 4).

**Fig. 5.**
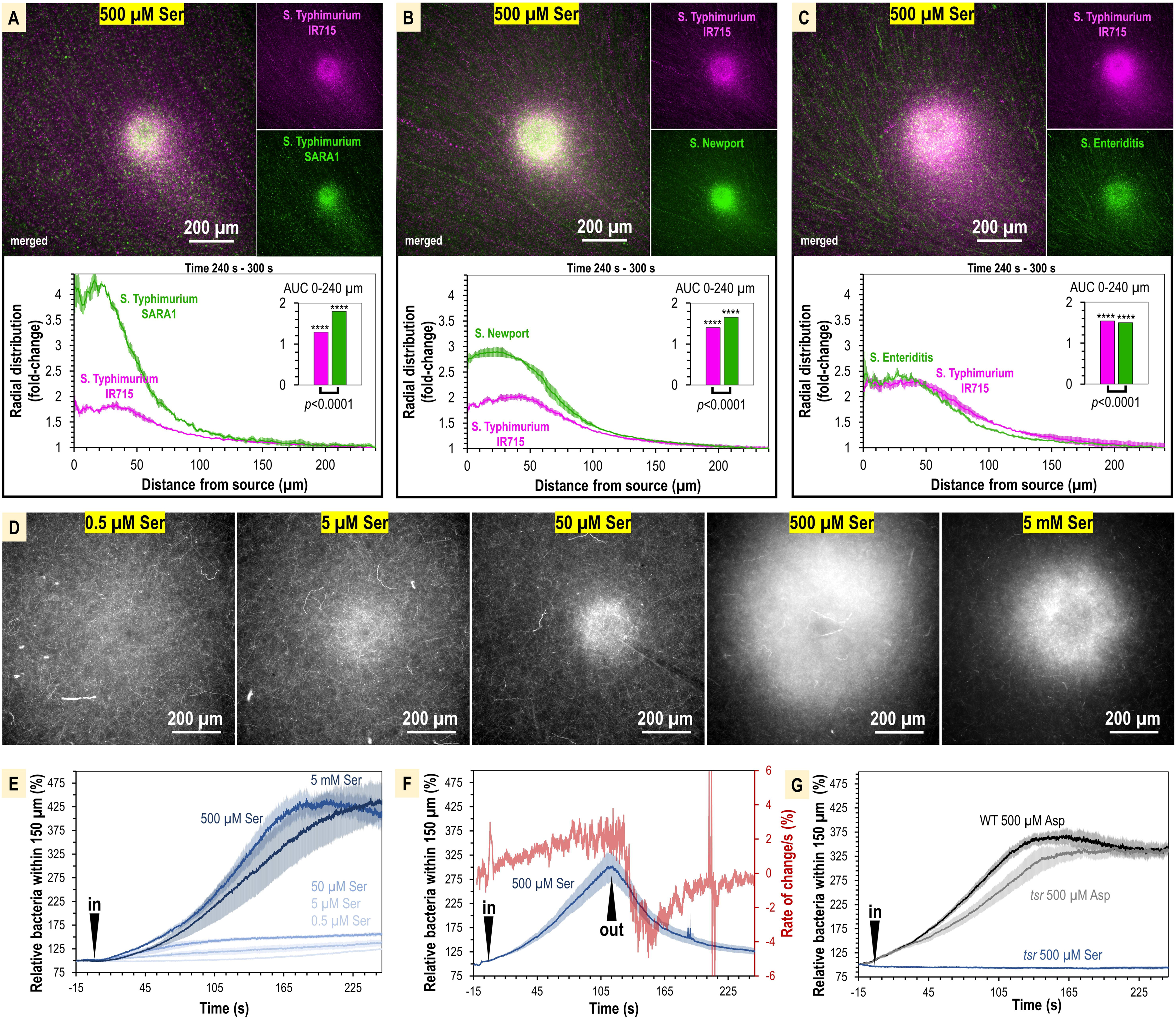
The concentration of L-serine in human serum is sufficient to mediate chemoattraction. A-C. CIRA competition experiments between *S.* Typhimurium IR715 (pink) and clinical isolates (green) in response to 500 µM L-serine (n=3, 37° C). D. Representative results showing max projections of *S.* Typhimurium IR715 at 240 – 300 s post CIRA treatment with L-serine concentrations (30° C). E. Quantification of multiple replicate experiments shown in D. F. Attraction and dispersion of *S.* Typhimurium IR715 following addition and removal of 500 µM L-serine source (30° C). G. *S.* Typhimurium IR715 WT or *tsr* mutant responses to L-aspartate or L-serine treatments (30° C).

We next used CIRA to test a range of L-serine sources spanning five orders of magnitude to define whether the concentrations of L-serine present in human serum are sufficient to drive a chemoattraction response. Within the 5 min timeframe of our experiments, the minimal source concentration of L-serine needed to induce chemoattraction is 0.5-5 µM. The [L-serine] source required for half-maximal chemoattraction (i.e., K_1/2_) is approximately 105 µM (Fig. 5D-E, Fig. S4). Based on our microgradient modeling, this corresponds to a local L-serine concentration of 1 nM for bacteria 100 µm from the source at *t*=300 s (Fig. S1, Fig. 4B). Therefore, we determined that the typical concentration of L-serine in human serum is >200-fold greater than the minimum required to elicit chemoattraction. To gain further insights into the dynamics of L-serine chemoattraction, we monitored chemotactic behavior in the presence of L-serine, and then removed the treatment (Fig. 5F). These experiments showed maximal attraction and dispersal rates to be similar, changing by approximately 4% per second (Fig. 5F). These findings emphasize the rapid dynamics through which chemotaxis can influence the localization of bacteria in response to microscopic gradients of chemoeffectors.

To substantiate our findings, we considered some alternative explanations for our data. First, we tested whether Tsr alone was required for chemoattraction to L-serine, or whether some other chemoreceptor, or form of taxis, such as energy or redox taxis, might contribute to the responses. However, we found when treated with L-serine, the *tsr* mutant showed no chemoattraction and behaved similarly to the chemotaxis-null *cheY* mutant (Fig. 5G). Second, we considered the possibility that the defect in serum attraction of the *tsr* mutant could be due to pleiotropic effects of the *tsr* deletion on chemotaxis signaling or motility, similar to the inhibited swimming motility of the *trg* mutant. We tested the ability of the *tsr* strain to respond to another chemoattractant, L-aspartate, which is sensed through the Tar chemoreceptor (Fig. 4A). We found that the *tsr* mutant mounted a robust chemoattraction response to 500 µM L-aspartate, similar in magnitude and rate to WT (Fig. 5G), supporting that chemotaxis to non-serine stimuli remains functional in the *tsr* strain. Along with the data showing the WT attraction to serum is diminished with serine racemase treatment, these results support that the mechanism of Tsr- mediated chemoattraction to serum is through direct recognition of L-serine.

### Serum provides a growth advantage for non-typhoidal S. enterica serovars

The robust serum attraction response conserved across diverse *Salmonella* serovars suggests serum, and L-serine present in serum, could be a source of nutrients during infection. Yet, serum also contains bactericidal factors that could inhibit bacterial growth ^29^. To address this uncertainty, we investigated whether active human serum provides a growth benefit for *Salmonella* when diluted into minimal media. In all strains surveyed, we found that serum addition enhances growth, and we saw no evidence of killing even for the highest serum concentrations tested (Fig. 6). The growth enhancement requires relatively little serum, as 2.5% v/v is sufficient to provide a 1.5-2.5-fold growth increase (Fig. 6).

**Fig. 6.**
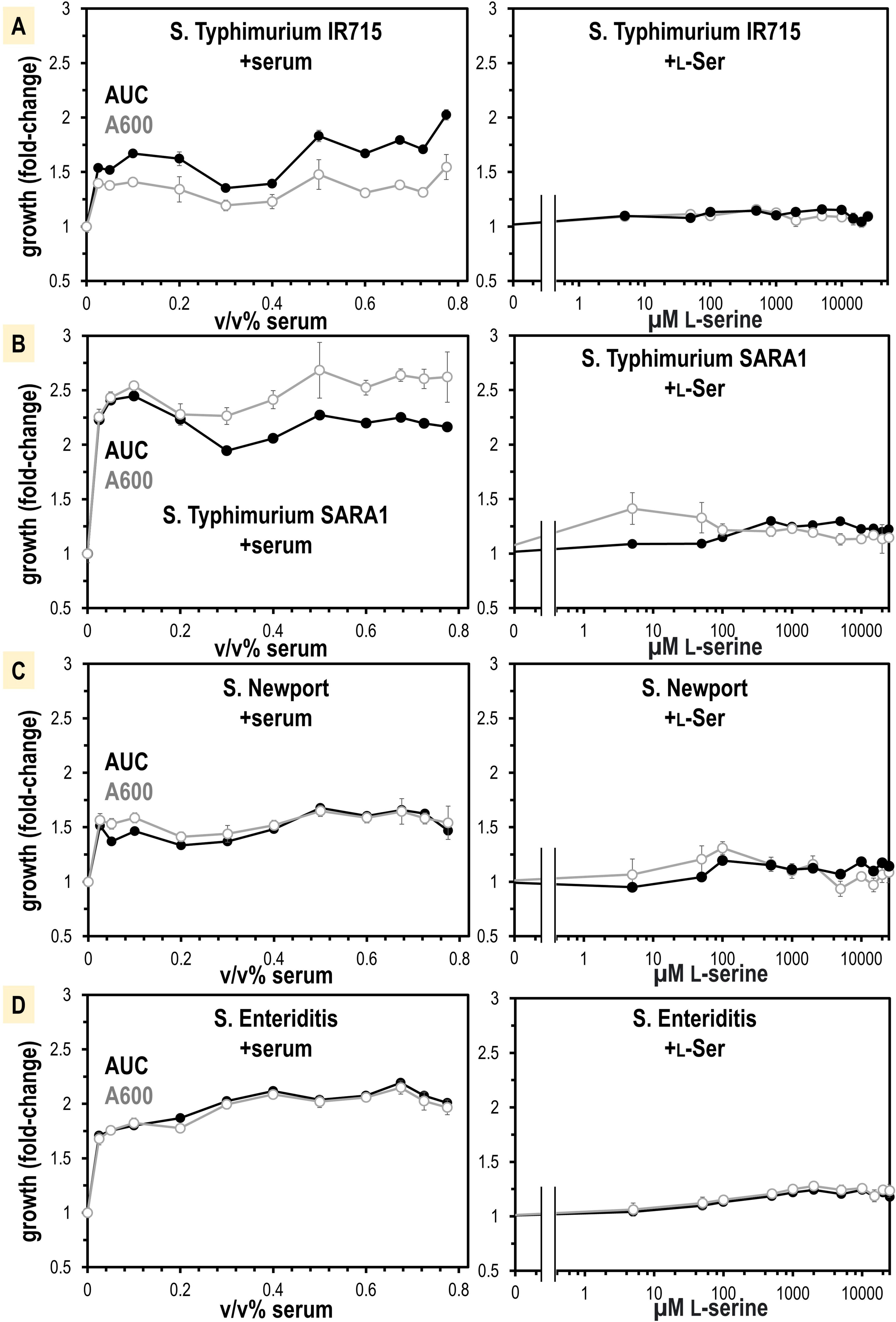
Non-typhoidal *S. enterica* strains obtain a growth benefit from human serum not recapitulated from L-Ser treatments alone. A-D. Growth is shown as area under the curve (AUC, black) and A600 at mid-log phase for the untreated replicates (gray, n=16). Data are means and error is SEM.

L-serine is not only a chemoattractant, but also an important nutrient for bacteria in the gut ^3^, so we hypothesized the growth benefit could be from L-serine. We determined by mass spectrometry that our human serum samples contain 241 µM +/- 48 total serine (L- and D- enantiomers), of which approximately 99% is expected to be L-serine ^44^ (Fig. S5). We attempted to treat human serum with a purified recombinant enzyme that degrades L-serine, serine dehydrogenase (SDS), to see whether serine-depleted serum would elicit less growth or chemoattraction. However, SDS treatments did not alter serum serine content so we abandoned this approach (Fig. S5). Instead, we performed a titration of purified L-serine and assessed its role in supporting a growth advantage. We found that only a very small benefit is achieved with the addition of L-serine, which did not recapitulate the larger growth benefit seen for serum addition (Fig. 6). This leads us to believe that L-serine functions as a molecular cue that directs *Salmonella* toward serum, but nutrients present in serum other than L-serine provide the major growth advantages.

### Structure of S. enterica Tsr in complex with L-serine

We next undertook structural studies to understand the specific recognition of L-serine by Tsr. The full-length Tsr protein includes a periplasmic ligand-binding domain (LBD), a transmembrane HAMP domain, and a cytosolic coiled-coil region, which oligomerizes to form trimers-of-dimers, and complexes with the downstream chemotaxis signaling components, CheA and CheW (Fig. 7A) ^45^. No experimentally-determined structure had been published for *S. enterica* Tsr (*Se*Tsr) and the single experimentally-determined Tsr structure to have captured the L-serine-binding interactions is a crystal structure of *Ec*Tsr LBD of moderate resolution (2.5 Å, PDB: 3ATP), in which the electron density for the ligand is weak and the orientation of the L-serine ligand is ambiguous ^46^. Despite the poorly-defined binding site interactions, this *Ec*Tsr crystal structure has guided numerous other studies of chemoreceptor signal transduction and nanoarray function ^45,47^.

**Fig. 7.**
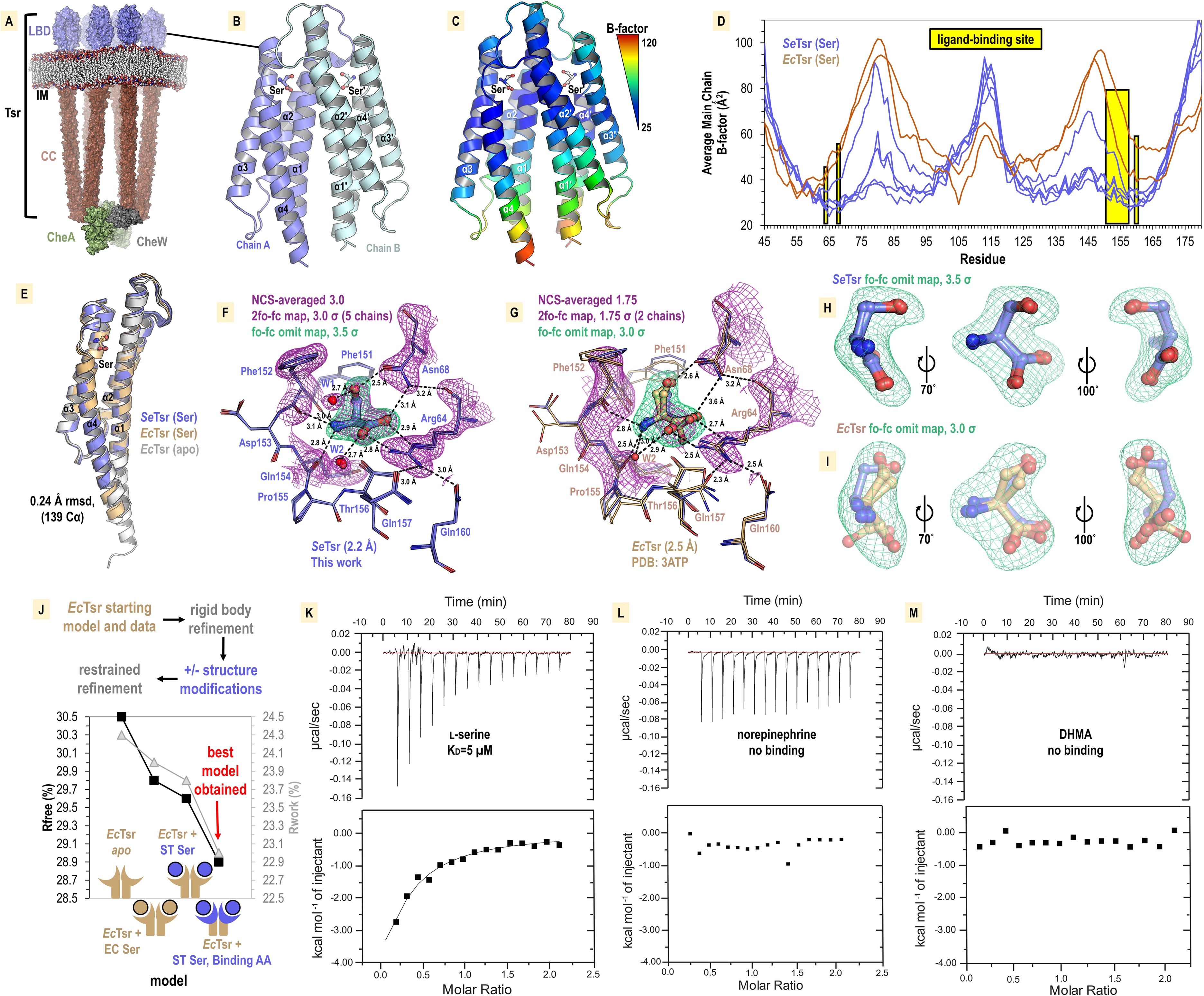
Structural mechanism underlying taxis to serum. A. Model of the core chemoreceptor signaling unit showing two full-length Tsr chemoreceptor trimer-of-dimers; coiled-coil region (CC), inner membrane (IM) ^45^. B. Crystal structure of *S. enterica* Tsr LBD dimer in complex with L-serine (2.2 Å). C-D. Relative order of the *Se*Tsr structure as indicated by B-factor (Å^2^). E. Overlay of chains from serine-bound *Se*Tsr (blue), serine-bound *Ec*Tsr (orange), and apo *Ec*Tsr (white). F. Binding of the L-serine ligand as seen with an overlay of the 5 unique chains of the asymmetric unit (AU) in the *Se*Tsr structure. Purple mesh represents averaged non- crystallographic symmetry 2f_o_-f_c_ omit map electron density (ligand not included in the density calculations). Green mesh represents f_o_-f_c_ omit map difference density for Chain A. Hydrogen bonds to the ligand are shown as dashed black lines with distances indicated in Angstroms (Å). G. The ligand-binding site of serine-bound *Ec*Tsr is shown as in F, with omit map f_o_-f_c_ electron density. The two chains of *Ec*Tsr in the AU are overlaid (orange) with one chain of serine-bound *Se*Tsr (blue). H-I. Closeup view of the L-serine ligand and f_o_-f_c_ omit map density for the *Se*Tsr (blue) and *Ec*Tsr (orange) structures, respectively. J. Paired refinement comparing resulting R- factors, as indicated. K-M. Isothermal titration calorimetry analyses of the *Se*Tsr LBD with L- serine, NE, or DHMA.

We recognized that in the prior *Ec*Tsr study, the methods described exchanging the protein crystal into a glycerol cryoprotectant. We hypothesized that during the glycerol soak serine leached out of the crystal leaving the binding site partially occupied and caused the electron density to be weak for the ligand and surrounding region. To capture a complex with a fully bound ligand site we grew crystals of the soluble periplasmic portion of the *Se*Tsr LBD with L-serine, at a high salt concentration that would serve as a cryoprotectant without further manipulation, and harvested crystals directly from drops to prevent leaching of the ligand from the binding site. After conducting extensive crystallization trials and examining x-ray diffraction data from over 100 crystals, we identified a small number of *Se*Tsr LBD crystals with sufficient quality for structure determination. The crystal that provided the highest-quality data (2.2 Å resolution, PDB: 8FYV) was grown in a mixture of buffers of pH 7.5-9.7 (refer to Methods, Table S2). To assess whether the basic pH had any impact on the protein structure, we solved a second structure from a crystal grown at pH 7-7.5 (Table S2, Fig. S6, PDB: 8VL8). By overlaying the two structures, it is apparent they are nearly identical, with no observable changes at the ligand-binding site (Fig. S6). Because the structure obtained at higher pH (PDB: 8FYV) exhibits superior quality and clearer electron density in critical regions of interest, it is this structure we refer to in subsequent sections.

The crystal structure of *Se*Tsr LBD contains five monomers in the asymmetric unit, providing five independent views of the L-serine binding site, with homodimers formed between Chains A and B, C and D, and E and its crystal symmetry mate, E′ (Fig. 7B). Lower B-factors in the *Se*Tsr ligand binding region are indicative of greater order, reflecting that our structure is fully serine-bound (Fig. 7C-D). *Se*Tsr and *Ec*Tsr possess high sequence similarity in the LBD region—100% identity of the ligand-binding residues and 82.1% identity over the entire periplasmic domain—and as expected, retain a similar global structure with all chains of serine- bound *Se*Tsr (5), serine-bound *Ec*Tsr (2), and apo *Ec*Tsr (2) overlaying within 0.24 Ǻ rmsd over 139 C_α_ (Fig. 7E).

### Molecular recognition of L-serine by Tsr

The higher resolution (+0.3 Ǻ) of the *Se*Tsr structure, and full occupancy of the ligand, provide a much-improved view of the interactions that facilitate specific recognition of L-serine (Fig. 7F). Using non-crystallographic symmetry averaging, we leveraged the five independent *Se*Tsr monomers to generate a well-defined 2f_o_-f_c_ map of the L-serine ligand and residues involved in ligand coordination (Fig. 7F, Movie 6). Omit-map difference density, which is calculated in the absence of a modeled ligand and reduces potential bias in the electron density map, was fit well by the placement of the L-serine ligand (Fig. 7F). In this orientation, the ligand is in an optimal energetic conformation that satisfies all possible hydrogen bonding interactions: the positively- charged peptide amine donates hydrogen bonds to the backbone carbonyl oxygens of Phe151, Phe152, and Gln154, the negatively-charged ligand carboxyl group donates hydrogen bonds to the Arg64 guanidinium group, the Asn68 side change amine, and a water (W2), and the ligand hydroxyl sidechain donates a hydrogen bond to the Asn68 sidechain oxygen, and accepts a hydrogen bond from a water (W1) (Fig. 7F, Movie 6). All five chains of the *Se*Tsr structure are consistent in the positions of the ligand and surrounding residues (Fig. 7F, Movie 6).

With the aid of the improved view provided by our *Se*Tsr structure, we noticed the L-serine is positioned differently than what was modeled into the weak density of the *Ec*Tsr structure (Fig. 7G-I, Movie 6). The *Ec*Tsr structure has the serine positioned with the sidechain hydroxyl facing into the pocket toward Asn68, and the orientations of Asn68, Phe152, Asp153, and Gln157 of the ligand binding pocket are modeled inconsistently, without justification, between the two *Ec*Tsr chains in the asymmetric unit (Fig. 7G-I). Calculating f_o_-f_c_ omit map density for both structures shows not only that our new *Se*Tsr ligand orientation fits the density of our structure well, but that it is a better fit for the data from the *Ec*Tsr structure (Fig. 7H-I). This is apparent by how the *Ec*Tsr serine C_β_ and sidechain hydroxyl are misaligned with the curvature of the f_o_-f_c_ map and that the carboxylate is partially outside of the electron density (Fig. 7I).

An analysis that can quantifiably determine which positions of the serine and ligand- binding residues result in the best model is to perform pairwise refinements and compare the crystallographic R_work_ and R_free_ statistics (Fig. 7J). Using as our starting model the *Ec*Tsr homodimer, and its deposited data from the protein databank, we performed refinements using identical strategies with three scenarios: no serine modeled in the binding site (*apo*), the published *Ec*Tsr serine pose, or the serine pose from our *Se*Tsr structure (Fig. 7J). The poorest resulting *Ec*Tsr model was *apo*, yielding R_work_/R_free_ values of +0.3%/+0.7% relative to the deposited model. This result supports the presence of the serine ligand in the structure. Substituting the serine with the pose observed in the *Se*Tsr structure led to an improved model, reducing the R_work_/R_free_ values by -0.2%/-0.2%. Lastly, we replaced both the serine ligand and the ligand-binding residues of the *Ec*Tsr model with those from our *Se*Tsr structure, which resulted in the highest quality model with meaningfully lower R_work_/R_free_ values of -1.0%/-0.9% (Fig. 7J). Consequently, we can conclude that the correct serine pose and ligand-binding site positions for both models are those we determined using the higher resolution *Se*Tsr structure, in which the L- serine side chain hydroxyl faces outward from the pocket, toward the solvent, and every hydrogen bonding group of the ligand is satisfied by the residues of the binding site (Fig. 7F, J).

### SeTsr LBD binds L-serine, but not NE or DHMA

Despite a lack of chemotactic responses to NE and DHMA in our CIRA experiments (Fig. S3, Movie 5, Movie S1, Data S1), our uncertainty lingered as to whether these neurotransmitters are sensed through Tsr. Prior work created theoretical models of NE and DHMA binding *Ec*Tsr at the L-serine site, and proposed this as the molecular mechanism underlying *E. coli* chemoattraction to these compounds ^42^. However, with our new experimentally-determined *Se*Tsr LBD crystal structure in hand, it is clear that the hydrogen bonding network is specific for L-serine (Fig. 7F, Movie 6), and we were doubtful such dissimilar ligands as NE or DHMA would be accommodated. We note that the amino acids that constitute the L-serine binding pocket are identical between *Se*Tsr and *Ec*Tsr (Fig. 7F-G), and so we reason that *Se*Tsr LBD and *Ec*Tsr LBD should have similar molecular functions and ligand specificity.

Despite several studies reporting a direct role of Tsr in directing chemoattractant responses to NE and DHMA, no direct evidence of ligand-binding has ever been presented. Thus, we performed isothermal titration calorimetry (ITC) experiments, a gold standard for measuring protein-ligand interactions, to study ligand recognition by *Se*Tsr LBD (Fig. 7K-M). As expected, L-serine produces a robust exothermic binding curve, exhibiting a K_D_ of approximately 5 µM (Fig. 7K). This matches well with our observation of *S. enterica* chemoattraction requiring a source of L-serine at least 0.5-5 µM (Fig. 5D-E) and with prior ITC data with *Ec*Tsr LBD, which also reported a 5 µM K_D_ ^46^. Conversely, neither NE nor DHMA showed any evidence of binding to *Se*Tsr LBD (Fig. 7L-M). These data confirm that *Se*Tsr is a receptor for L-serine, and not NE or DHMA, and further support the molecular mechanism of Tsr-mediated serum attraction to be through L-serine.

### Relevance of bacterial taxis to serum in other systems

Our chemotaxis experiments with the *tsr* strain show that while Tsr is not the only chemoreceptor involved in taxis to serum for *S. enterica*, it does play an important role through chemoattraction to the high concentration of L-serine within serum. We hypothesized that attraction to serum, mediated through Tsr, might extend beyond *S. enterica* to other bacterial species. The biological distribution of Tsr chemoreceptors has not been previously characterized, so we applied our structural insights to define a motif consisting of the amino acids involved in L-serine recognition. When we analyzed the genomes of Enterobacteriaceae genera such as *Escherichia*, *Citrobacter*, and *Enterobacter* we found their Tsr orthologues to possess this motif (Fig. 8A). We next conducted a comprehensive search for chemoreceptor genes containing the L- serine recognition motif, creating a database containing all organisms that possess putative Tsr orthologues (Fig. 8B, Data S2). The biological distribution of Tsr reveals the chemoreceptor to be widely conserved among Gammaproteobacteria, particularly within the families Enterobacteriaceae, Morganelliaceae, and Yersiniaceae (Fig. 8B). We discovered that many WHO priority pathogens are among the species that have Tsr, leading us to suspect these pathogens and pathobionts also perform serum taxis.

**Fig. 8.**
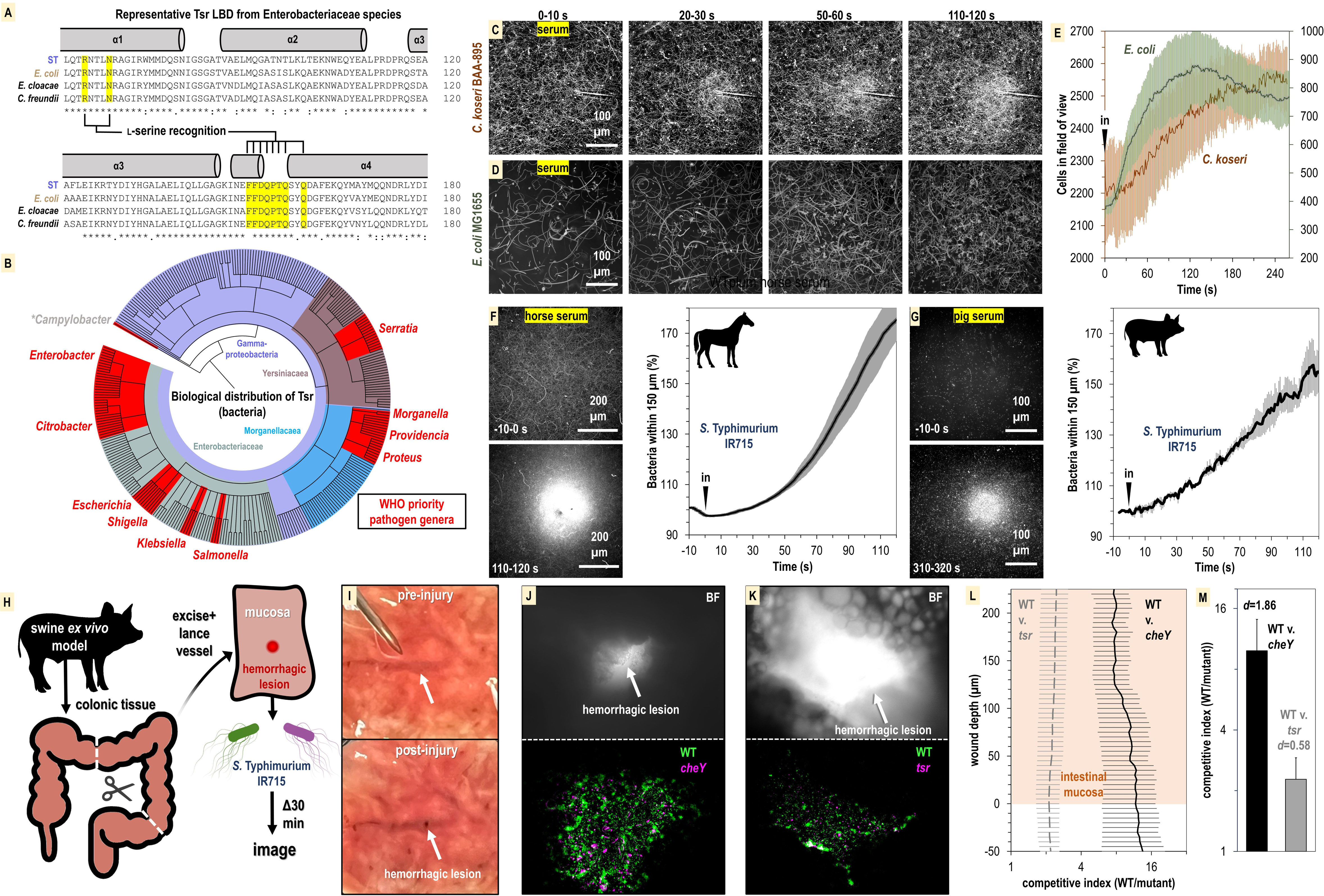
Relevance of bacterial vampirism in other systems. A. Sequence conservation of the Tsr ligand-binding domain among representative Enterobacteriaceae orthologues with residues of the L-serine ligand-binding pocket highlighted. B. Biological distribution of Tsr orthologues. WHO priority pathogen genera are highlighted in red. C-D. Response of *C. koseri* BAA-895 (n=3, 37° C) and *E. coli* MG1655 to human serum (n=3, 30° C), shown as max projections at indicated time points. E. Quantification of chemotaxis responses. Plotted data shown are shown on separate y-axes as mean cells in the field of view averaged over 1 s. F-G. Response of WT *S.* Typhimurium IR715 to horse serum or pig serum. H. Overview of swine *ex vivo* colonic hemorrhagic lesion model. I. Representative images of colonic mucosa before (top) and after (bottom) lancing of vasculature. J-K. Representative images showing bacterial localization into colonic hemorrhagic lesions in co-inoculation experiments with WT (*gfp*) and *cheY* (*dtom*), or WT (*gfp*) and *tsr* (*dtom*) S. Typhimurium IR715. The lesion is shown in brightfield (BF, top), and gfp/dtom fluorescent channels (bottom). L. Competitive indices of migration into lesions. The mucosa is set to a y-axis value of 0, and migration further into the lesion is reflected by increasing y-axis values. M. AUC quantification of bacterial lesions localization shown in panel L (0-225 µM). Effect size (Cohen’s *d*) is noted. Data shown are means and error bars show SEM.

To address this question, we performed CIRA analyses with human serum and the bacterial strains *E. coli* MG1655 and *C. koseri* BAA-895, the latter being a clinical isolate with previously uncharacterized chemotactic behavior. We find that both these strains exhibit attraction to human serum on time scales and magnitudes similar to experiments with *Salmonella* (Fig. 8C-E, Movies S2-S3). Without further genetic analyses in these strain backgrounds, the evidence for Tsr mediating serum taxis for these bacteria remains circumstantial. Nevertheless, taxis to serum appears to be a behavior shared by diverse Enterobacteriaceae species and perhaps also Gammaproteobacteria priority pathogen genera that possess Tsr such as *Serratia*, *Providencia*, *Morganella*, and *Proteus* (Fig. 8B).

So far, our focus has been on bacterial taxis towards human serum. However, given the presence of L-serine in the serum of various animal species, we were interested in exploring if serum from other mammals could also attract bacteria. We prepared horse serum from whole horse blood through clotting and centrifugation to use in CIRA experiments with *S.* Typhimurium IR715. Similar to human serum, the bacterial population rapidly swam toward the horse serum source, with the relative number of cells in close proximity to the source nearly doubling within 120 seconds of exposure (Fig. 8F). To further investigate this concept, we utilized serum we collected from a third species, a single domestic pig. The data revealed that pig serum similarly induces a rapid attraction of *S.* Typhimurium (Fig. 8G). This suggests that the phenomenon of serum attraction is relevant to diverse host-microbe systems and is not specific to bacteria associated with humans. Considering the data from all CIRA experiments, while some differences are observed in strain responses to serum sources (*S. enterica* serovars, *E. coli*, and *C. koseri*) and serum from different animals (human, horse, and pig), all WT strains examined share the behavior of rapid taxis towards serum.

As a preliminary investigation into whether serum taxis is involved in bacterial pathogenesis, we developed an enterohemorrhagic lesion model to investigate if chemotaxis and Tsr can mediate bloodstream entry. We found that pig serum stimulates pathogen attraction (Fig. 8G), and swine recapitulate many aspects of human gastrointestinal physiology ^48^, so we utilized swine colonic tissue from the same animal for *ex vivo* experimentation (Fig. 8H). We created an enterohemorrhagic lesion by a single lance of the tissue vasculature from the mucosal side, and then exposed the wound to a 1:1 mixture of motile *S.* Typhimurium IR715 WT and chemotaxis mutant (Fig. 8H-I). Following 30 minutes of exposure, the localization of the bacteria in proximity to the lesion was determined through fluorescence microscopy and enumerated as a competitive index of WT vs. mutant. By imaging through the z-plane in five-micron intervals we were also able to quantify bacterial colonization as a function of wound depth. In the context of this *ex vivo* model, we observed that the bacteria were readily able to penetrate >200 µm into the wounded vasculature (Fig. 8J-M). Strikingly, the WT efficiently localizes toward, and migrates into, the hemorrhagic lesion, outcompeting the *cheY* and *tsr* mutants by approximately 10:1 and 2:1 (Fig. 8J-M). These results align with our other data indicating that serum taxis requires CheY, and that Tsr contributes, but is not the only chemoreceptor involved in mediating attraction. Additional investigation is required to confirm the occurrence of these behaviors during infection, but it seems plausible that bacterial taxis toward sources of serum in the host gut environment occurs, potentially contributing to the development of bacteremia.

## Discussion

The most common outcome for gastrointestinal bacterial infections is that they are resolved by the immune system without chronic infection or lingering pathologies (Fig. 1). To understand why, in rare circumstances, infections do persist, become chronic, systemic, and life-threatening, we must uncover the mechanisms at the crossroads where pathogens divert from the routine course of infection and do something unusual. An emerging area of interest in the axis of gut- microbe health is how the behavior of gut microbes can shift in response to inflammation, antibiotic usage, and pathogen invasion, into a state of dysbiosis recalcitrant to treatment ^15,49–51^. We can learn about the factors that stimulate microbial-induced pathologies by studying how bacteria respond to host-derived stimuli that are unique to the diseased gut ^51^. Here, we have investigated how bacterial chemosensing functions to respond to a source of serum, a chemical feature enteric bacteria only encounter in the event of GI bleeding.

We demonstrated that serum from humans and other animals contains components that act as chemoattractants and nutrients for Enterobacteriaceae known as instigators of enteric bleeding and causal agents of bacteremia and sepsis (Fig. 2-3, Fig. 6, Fig. 8). Bacterial infiltration of wounded enteric vasculature, mediated by chemotaxis and the chemoreceptor Tsr, suggests serum attraction plays a role in the bloodstream entry of these bacterial species. In a broader context, the attraction of Enterobacteriaceae to serum aligns with an emerging understanding of how bacterial chemotaxis can drive tropism for sites of damaged tissue, injury, and inflammation (Fig. 1) ^3,23,52–56^. This phenomenon of bacterial attraction to serum through chemotaxis to access serum nutrients represents a novel pathogenesis strategy, which we term ‘bacterial vampirism’ (Fig. 9).

**Fig. 9.**
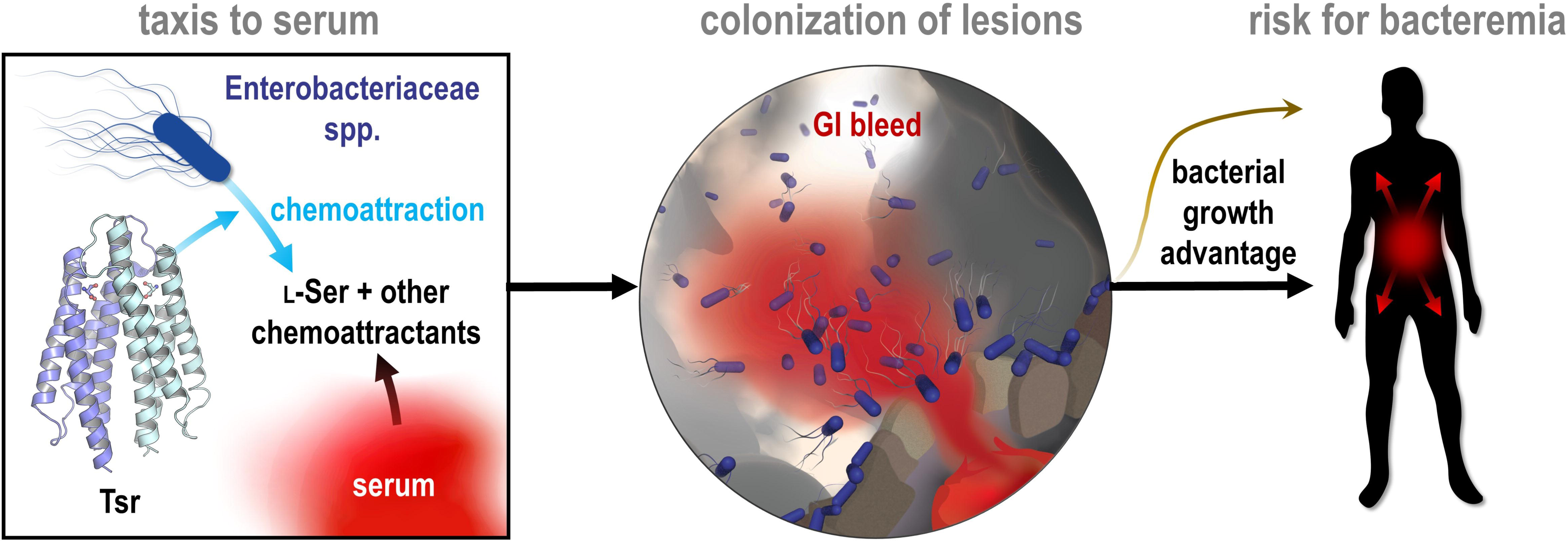
Model of bacterial vampirism. Serum contains high concentrations of L-Ser, and other chemoattractants, that are recognized by chemoreceptors, including Tsr, to drive Enterobacteriaceae taxis toward serum. Taxis to serum promotes colonization of enterohemorrhagic lesions and provides a bacterial growth advantage from nutrients acquired from serum. The bacterial behavior of seeking and feeding on serum, which we have defined as “bacterial vampirism” may represent a risk factor for bacterial entry into the bloodstream.

### A new ecological context for L-serine chemoattraction

We show here that the bacterial attraction response to serum is robust and rapid; the motile population reorganizes within 1-2 minutes of serum exposure to be significantly biased toward the serum source (Fig. 2, Fig. 8). Serum taxis occurs through the cooperative action of multiple bacterial chemoreceptors that perceive several chemoattractant stimuli within serum, one of these being the chemoreceptor Tsr through recognition of L-serine (Fig. 4). While Tsr is known as a bacterial sensor of L-serine ^3,45,57^, the physiological sources of L-serine in the gut sensed by Tsr have not been well-defined. Although a major source is dietary, recent research has suggested that L-serine from damaged tissue may play a significant role in the diseased gut environment and drive opportunistic pathogenesis. In a study using a dextran sodium sulfate (DSS)-colitis model to investigate L-serine utilization by Enterobacteriaceae, L-serine was identified as a critical nutrient providing metabolic and growth advantages in the inflamed gut ^51^. Moreover, colitis was found to stimulate substantial increases in the luminal availability of most amino acids, with the average serine content nearly doubling ^51^. A notable characteristic of the DSS- colitis model is intestinal bleeding ^58^, suggesting that the enrichment of amino acids, including serine, could be attributed to the leakage of host metabolites from serum. The ability of pathogenic Enterobacteriaceae to exploit growth advantages from L-serine during colitis may be linked to their highly sensitive and rapid chemoattraction to this effector emitted from human serum.

Our structural and molecular investigations into *S. enterica* Tsr affirm its role as a serine sensor, revealing a highly optimized and selective ligand-binding site for the recognition of L- serine (Fig. 7, Movie 6). Although the host-derived metabolites NE and DHMA have been reported to be chemoattractants for *E. coli* sensed directly through Tsr, we were unable to corroborate these results in *S.* Typhimurium in chemotaxis experiments, nor binding assays with the purified recombinant LBD, despite the high sequence conservation between the two orthologues (Fig. S3, Fig. 7L-M, Fig. 8A, Data S1). Our inability to substantiate a structure- function relationship for NE/DHMA signaling indicates these neurotransmitters are not ligands of Tsr, or, at the very least, are not broadly relevant as chemoeffectors sensed through Tsr orthologues. In contrast, we saw that L-serine drove chemoattraction to a source as low as 500 nM, all strains tested were attracted to L-Ser, a binding K_D_ to *Se*Tsr LBD of 5 µM was established through ITC, and was captured in complex in the *Se*Tsr LBD crystal structure with clearly-defined electron density (Fig. 5, Fig. 7). Our study is not the only one to raise questions regarding the physiological relevance of NE/DHMA as chemoattractants. A study examining *E. coli* sensing of NE/DHMA by Tsr showed little chemoattraction occurs in the range of 10-100 µM, and at higher (millimolar) concentrations a chemorepulsion response was observed ^59^. Notably, the concentrations required in the latter study to observe a chemotactic response are far above what is thought to be physiologically relevant for tissue or serum ^60^. Given that treatment of serum with serine racemase diminishes taxis toward serum and eliminates the WT taxis advantage over *tsr* (Fig. S3), our data support that L-serine is the chemoattractant sensed through Tsr that confers serum attraction.

An unanticipated result from obtaining the first structures of *Se*Tsr LBD (PDB: 8FYV) is that the well-resolved ligand-binding site contradicts key interactions as they were modeled in the earlier *Ec*Tsr LBD structure (3ATP), apparently owing to the difficulty of interpreting the weak electron density (Fig. 7F-I). Through comparative refinements, we show that changing the ligand position and binding site residues of the *Ec*Tsr LBD model to be as they are in the *Se*Tsr LBD structure demonstrably improves the model (Fig. 7J). Therefore, the differences in the models are not attributable to them being two different proteins, rather the *Ec*Tsr LBD structure is modeled incorrectly, and both structures should be modeled with the serine pose and ligand- binding site as they are in the *Se*Tsr LBD structure (Fig. 7F). The *Ec*Tsr LBD structure has informed the creation of full-length atomic models of Tsr, and the core signaling unit, for use in molecular simulations and fitting of cryo-EM data ^45,47,61^. Since we now have in hand a higher resolution and better-resolved Tsr crystal structure, future studies of this sort should benefit from the improved understanding of L-serine recognition. The *Se*Tsr LBD structure was also helpful in accurately defining the signature motif of Tsr orthologues, which allowed us to mine genomic databases and report the first extensive characterization of which bacterial species possess this chemoreceptor (Fig. 8A-B). From this database of Tsr sequences we were able to visualize the distribution of Tsr orthologues across biology, which strikingly are enriched among Enterobacteriaceae, but are also present in several other families that include WHO priority pathogens (Fig. 8B, Data S2). This analysis confirms that the bacterial species most commonly associated with bacteremia in patients with IBD possess the Tsr chemoreceptor (Fig. 8B, Data S2) ^25^.

In the context of *Salmonella* pathogenesis, it is interesting to note that diverse serovars, which vary in terms of host specificity and epidemiology, exhibit serum attraction and have the ability to utilize serum as a nutrient (Fig. 3, Fig. 6). The specific nutrients responsible for the growth benefits derived from serum remain undefined, however, one potential candidate is the presence of energy-rich glycoconjugates, which were previously implicated as nutrients for bacteria within the inflamed intestine based on the upregulation of galactose utilization operons and the prevalence of lectin-positive stained tissue ^62^. Enteric Peyer’s patches are primary invasion sites for non-typhoidal *Salmonella*, and these structures are situated close to vasculature that can be damaged through the pathogen’s destruction of microfold (M) cells, causing localized bleeding ^63^. *S. enterica* Typhimurium uses the chemoreceptor Tsr to locate and invade Peyer’s patches ^32^, which could involve L-serine chemoattraction originating from serum or necrotic cells. Although GI bleeding is relatively uncommon in *Salmonella* infections overall, it afflicts 60% of infected children under the age of five ^64^. Therefore, while we acknowledge that serum attraction is not a routine pathogenesis strategy employed by *Salmonella*, the significant number of bacterial-induced GI bleeding cases, and the association between GI bleeding and bacterial invasion into the bloodstream, provide opportunities for bacterial vampirism to be involved in infection outcomes (Fig. 9) ^24,27,65^.

## Materials and Methods

### RESOURCE AVAILABILITY

#### Lead contact

Further information and requests for resources and reagents should be directed to and will be fulfilled by the lead contact, Arden Baylink.

#### Materials availability

Strains and plasmids generated in this study will be made available upon request by the Lead Contact with a completed Materials Transfer Agreement.

#### Data availability

The crystal structures of L-serine-bound *Se*Tsr LBD are deposited to the protein databank as PDB: 8FYV and 8VL8. Any additional information required to reanalyze the data reported in this paper is available from the Lead Contact upon request.

### EXPERIMENTAL MODEL AND SUBJECT DETAILS

#### Bacterial strains

Strains of *S. enterica*, *E. coli*, and *C. koseri* used in this study are described in Table S1. To generate fluorescent strains, electrically-competent bacterial cultures were prepared through successive washing of cells with ice-cold 10% glycerol, and then transformed by electroporation (Biorad GenePulser Xcell) with pxS vectors containing either sfGFP, or mPlum, and AmpR genes ^66^. Transformants were isolated through growth on selective media and stored as glycerol stocks for subsequent use. To prepare motile bacterial cells for CIRA experiments, bacterial cultures were grown shaking overnight in 2-5 ml tryptone broth (TB) and 50 µg/ml ampicillin (TB+Amp) at 30° C or 37° C, as needed. The following day, 25 µl of overnight culture was used to inoculated 25 ml of fresh TB+Amp and grown shaking for 3-5 hours at the desired temperature to reach A_600_ of approximately 0.5. To better isolate responses to effectors of interest and remove confounding variables, such as pH variation and quorum-signaling molecules, we washed and exchanged the cells into a buffer of defined composition. Bacterial cultures were centrifuged at 1,500 g for 20 minutes and exchanged into a chemotaxis buffer (CB) containing 10 mM potassium phosphate (pH 7), 10 mM sodium lactate, and 100 µM EDTA. Cultures were diluted to approximately A_600_ 0.2 (or as indicated in figure legends) and allowed to recover rocking gently for 30-60 minutes to become fully motile.

For *in vitro* growth analyses, *S. enterica* strains were grown in Luria-Bertani (LB) media shaking overnight at 37° C. The following day, cultures were pelleted by centrifugation and resuspended in a minimal media (MM) containing 47 mM Na_2_HPO_4_, 22 mM KH_2_PO_4_, 8 mM NaCl, 2mM MgSO_4_, 0.4% glucose (w/v) 11.35 mM (NH_4_)_2_SO_4_, 100 μM CaCl_2_. 5 µl of the overnight cultures at 0.05 A_600_ were used to inoculate fresh solutions of 200 µl of MM, or additives diluted into MM (human serum or L-serine), in a 96-well microtiter plate. A plate reader was used to monitor cell growth while shaking at 37° via A_600_ readings every 5 minutes.

### METHOD DETAILS

#### Chemosensory injection rig assay (CIRA)

Our CIRA system is based on prior work, with several notable changes to methodology, as described ^5,6,67^. The CIRA apparatus was constructed using a pump for injection (either a refurbished Eppendorf Transjector 5246 or Femtojet 4i) and universal capillary holder, with solutions injected through Femtotip II glass microcapillaries (Eppendorf), and the microcapillary position controlled with a MP-285 micromanipulator (Sutter) at a 30° angle of attack. To generate a microgradient, a constant flow from the microcapillary was induced by applying compensation pressure (P_c_) of 35 hPa, unless otherwise specified. The stability of the microgradient under these conditions was determined with Alexa488 dye (ThermoFisher), and the flow was determined empirically with methylene blue dye to be approximately 300 femtoliters per minute at 30° C (Fig. S1). Pooled off-the-clot human serum was obtained from Innovative Research; human serum was deidentified and had no additional chemical additives and is presumed to be fully active. Horse serum was prepared from whole horse blood from Innovative Research. Pig serum was prepared from whole blood collected from a single animal. Treatment solutions were filtered through a 0.2 µm filter and injected as-is (serum) or diluted in CB (serine, aspartate, NE, DHMA). Serine racemase treatment of human serum was performed with the addition of 5 µl of proprietary serine racemase solution from a DL-serine assay kit (Abcam) to 1 ml of serum for 3 h. For each CIRA experiment, a fresh pond of 50 µl of motile bacteria was mounted open to air on a 10-well slide (MP Biomedicals), and the microcapillary containing the treatment of interest was lowered into the pond. Bacterial responses were imaged with an inverted Nikon Ti2 Eclipse microscope with an enclosed heated sample chamber. Temperature of experiments were at 37 °C, unless otherwise indicated.

#### CIRA microgradient modeling

Diffusion is modeled as a 3D process in which the diffusive species is slowly and continuously introduced at a fixed point in a large ambient fluid volume. The species to be injected is prepared at concentration *M_s_* (typically in the range of 0.5 µM-5 mM) and is injected at a volume rate Q = 305.5 fl/min. The species diffuses into the ambient fluid with diffusion constant D. The governing equation for the diffusion of a species introduced continuously at a point source is:

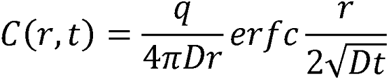

where *r* is the distance from the point source, *t*, is the time from injection initiation, and *q = M_s_Q* is the injection rate of the species, and *C* is the species concentration. We can simplify the presentation by defining a characteristic length scale *r*_0_,“, characteristic time *t*_0_,”, and dimensionless variables as:

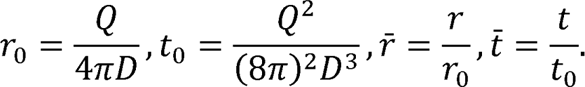

Then, we have the diffusion-driven concentration model:

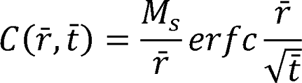

Representative diffusion coefficients are: A488, 4.00×10^−6^cm^2^/s; L-serine, 8.71 × 10^−6^cm^2^/s; L-aspartate, 9.35×10^−6^cm^6^/s. Due to the small injection rate, our assumption of a point source leads to a model that is valid at distances, *r* ≫ *r*_0_ ∼ nm and times *t* ≫ *t*_0_ ∼ 1 ns. We also consider the total species quantity integrated along a viewing direction. The result is:

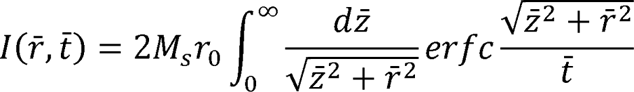

where 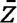 is the unitless integration variable along the viewing direction. This integral was evaluated numerically by considering the sequence of points 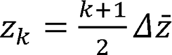for , - 1,2,3,… and,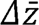 some small interval. Then, we have the discrete integral approximation:

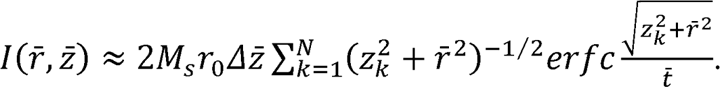

Computations for this work used 1 µm steps extending out to *r* = 500 μm. That is,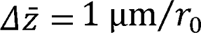, and .*N* = 500.

#### Ex vivo swine hemorrhagic lesion model

Swine intestinal tissue was procured from a single 8-week-old farm-reared animal (the same animal from which serum was drawn), in accordance with Washington State University Institutional Animal Care and Use Committee and Institutional Biosafety Committee approval. Intact colonic tissue was excised, and an incision was made along its length to flatten the tissue and expose the mucosa. Sections of tissue approximately 2.5 square centimeters in size were prepared for experimentation through gentle washing with saline solution, followed by CB, to remove fecal contents and debris. A sterile Gawal Sharp Point was used to lance the vasculature to generate a model enterohemorrhagic lesion. The lesion site was positioned in a MatTek dish over a pond of 50 µl of motile bacterial cells (1:1 fluorescently-tagged WT and mutant in CB, at a total A_600_ of 1.0). After 30 minutes, bacterial localization into the lesion was visualized using an inverted Nikon Ti2 Eclipse microscope. Images were captured at 20x magnification proceeding through the Z-plane in 5 µm intervals.

#### Cloning and recombinant protein expression

Cloning of the *S. enterica* Tsr LBD construct for recombinant protein expression was performed as a service by Genscript Biotech Corp. The sequence of the periplasmic portion of the ligand- binding domain of *S. enterica* Typhimurium Tsr (gene *STM4533*), corresponding to residues 32- 187 of the full-length protein, was encoded into a pet-30a(+) vector (Tsr-LBD-pet-30a(+)), at the NdeI and HindIII sites, with a short N-terminal TEV cleavage tag (MENLYFQ) such that the final expressed protein sequence was:

MENLYFQSLKNDKENFTVLQTIRQQQSALNATWVELLQTRNTLNRAGIRWMMDQSNIG SGATVAELMQGATNTLKLTEKNWEQYEALPRDPRQSEAAFLEIKRTYDIYHGALAELIQ LLGAGKINEFFDQPTQSYQDAFEKQYMAYMQQNDRLYDIAVEDNNS

Chemically-competent Rosetta BL21(DE3) *E. coli* (Millipore Sigma) were transformed by heat shock with the Tsr-LBD-pet-30a(+) vector, and transformants identified by growth on selective media containing 20 µg/ml (LB+Kan). Cells were grown overnight in 5 ml LB+Kan. The following day, 1 ml of overnight culture was used to inoculate 1 L of fresh LB+Kan, and cultures were grown to OD_600_ 0.6-0.8 and induced with 0.4 mM isopropyl β-D-1-thiogalactopyranoside (IPTG). After growth for 3 H at 37° C cells, were harvested by centrifugation.

#### Purification of recombinant SeTsr LBD

The cell pellet was resuspended in a lysis buffer containing 50 mM Tris pH 7.5, 0.1 mM DTT, 1 mM EDTA, 5 mg DNAse I, and 1 cOmplete protease inhibitor tablet per 1 L of culture (Sigma Aldrich), and cells were lysed by sonication. After, the lysate was kept on ice and adjusted to 20 % ammonium sulfate saturation and stirred at 4° C for 30 minutes. Lysate was centrifuged at 15,000 rpm for 30 minutes in a Beckman ultracentrifuge. The soluble fraction was retained, and an ammonium precipitation trial was conducted; the 20-40% fraction contained the majority of the Tsr LBD protein and was used for subsequent purification. The protein solution was dialyzed for 16 hours against 4 L of 20 mM Tris, pH 7.5, 20 mM NaCl, and 0.1 mM EDTA, then run over an anion exchange column and FPLC (Akta). The purest fractions were pooled and treated with 0.3 mg/ml TEV protease, and the protein solution was dialyzed against 4 L of 50 mM Tris pH 8, 0.5 mM EDTA, and 1 mM EDTA for 48 h at 4° C. Subsequently, the cleaved protein solution was exchanged into a buffer of 50 mM Tris pH 7.5, 1 mM EDTA, and 150mM NaCl, and purified by gel filtration with an S200 column and FPLC. Pure protein fractions were pooled, concentrated to 7 mg/ml, and flash frozen in liquid N_2_.

#### Protein crystallography

Initial crystallization trials of *Se*Tsr LBD were performed with either TEV-cleaved or uncleaved protein samples at 7 mg/ml with +/- 2 mM L-serine and using 96-well matrix screens set up with a Mosquito robot (SPT Labtech). We only observed crystal hits with the cleaved, serine-treated crystals, the best of which was 0.056 M sodium phosphate, 1.344 M potassium phosphate, pH 8.2. This condition was further optimized to be 1.5 µl *Se*Tsr LBD protein (7 mg/ml), 0.5 µl of 8 mM L-serine, and 1.5 µl 1.69 M potassium phosphate pH 9.7, grown via hanging drop vapor diffusion with a 1 ml reservoir of 1.69 M potassium phosphate pH 9.7 at 22° C. Crystals were scooped directly from drops and flash frozen in liquid N_2_. X-ray diffraction data were collected at the Berkeley Advanced Light Source (ALS) beamline 5.0.3. Out of over 100 crystals examined, only one diffracted to high resolution and was not impacted by crystal twinning. Data were indexed with DIALS ^68^, scaled with Aimless ^69^, and found to correspond well to space group C2_1_. A conservative final resolution cutoff of 2.2 Å was applied on the basis of CC_1/2_ >0.3 and completeness >50% in the highest resolution shell ^70^.

The serine-bound *Ec*Tsr structure (PDB: 3ATP) was utilized as a molecular replacement search model with Phaser-MR in Phenix ^71^ to solve the *Se*Tsr LBD dataset with five monomers in the asymmetric unit. 10% of the data were designated as R_free_ flags and the initial model was adjusted by setting all B-factors to 30 Å^2^ and coordinates were randomized by 0.05 Å to reduce bias from the starting model. Subsequent model building with Coot ^72^ and refinement with Phenix enabled placement of residues 42-182 and the serine ligand. However, residues 32-41 and 183-187 could not be resolved and were not modeled, causing the R/R_free_ values to be elevated for a model of this resolution. Riding hydrogen atoms and translation, libration, screw refinement were applied and reduced R factors. The strongest remaining difference peak is along a symmetry axis. The final model R/R_free_ values were 24.1/25.8 %, with a 99^th^ percentile all-atom MolProbity clashscore, and a 100^th^ percentile overall MolProbity score ^73^.

Because the protein was in a buffer at pH 7.5, but the crystallization solution was at 9.7, there was uncertainty about the true pH of the crystalline sample and how this might impact ligand binding interactions. As a control, we crystallized the protein in a mother liquor of 1.62 M potassium phosphate pH 7, with 0.5 µl of 3 mM L-serine, at 22° C. Solution of this structure revealed no remarkable differences in the ligand binding interactions or in the global structure (Fig. S6). The data quality suffered from ice rings, resulting in maps with high noise, and so we restricted refinement to a conservative 2.5 Ǻ cutoff. Crystallographic statistics for these structures are listed in Table S2. The pH 7.5-9.7 and pH 7-7.5 structures were deposited to the protein data bank as entries 8FYV and 8VL8, respectively. On the basis of the higher quality maps, and stronger and more clearly interpreted electron density, we recommend structure 8FYV for use in future structural studies of *Se*Tsr.

For comparisons of model quality between *Se*Tsr LBD (8FYV) with *Ec*Tsr (3ATP), we re-refined 3ATP with its deposited data using rigid body refinement to obtain a starting model for subsequent evaluation. Then, we performed a series of restrained refinements with Phenix using identical R_free_ flags and identical strategy of five cycles of xyz reciprocal space, xyz real space, individual isotropic B-factor, adp weight optimization, and stereochemistry weight optimization. For these refinements we used the *Ec*Tsr LBD structure +/- the following modifications: (1) the deposited L-Ser present, (2) the deposited L-Ser removed (*apo*), (3) the L- Ser replaced by the *Se*Tsr L-Ser ligand, or (4) both the L-Ser and the ligand binding site residues replaced by those from the *Se*Tsr LBD structure. These comparisons show the best model, as evidenced by meaningfully reduced R_work_ and R_free_ values, is obtained with the position of the L- Ser and ligand binding residues from the higher resolved *Se*Tsr LBD structure (Fig. 7J).

#### Isothermal titration calorimetry ligand binding studies (ITC)

ITC experiments were performed in 50 mM Tris, 150 mM NaCl, 1 mM EDTA, pH 7.5, at 25 °C. We dialyzed the protein into the experimental buffer and dissolved the small molecule ligands into the same buffer. Protein concentrations were ∼50 μM; titrant concentrations were 500 μM. Samples were degassed prior to experiments. All experiments were performed on a MicroCal ITC-200 system (GE Healthcare), with the gain set to “low” and a syringe stir speed of 750 rpm. Titration data for the serine experiments were fit to a single-site binding model using the built-in ITC analysis software.

#### Mass Spectrometry

Determination of molar content of total serine in human serum samples was performed as a service through the University of Washington Mass Spectrometry Center. Samples were analyzed on the Waters TQ #1 instrument using a Thermo Hypersil Gold PFP column (2.1 x 100) with 0.1% heptafluorobutyric acid (HFBA) in water and acetonitrile.

### QUANTIFICATION AND STATISTICAL ANALYSIS

#### Quantification of CIRA data

To determine relative numbers of cells over time, a ratio of fluorescence intensity per cell was calculated using ImageJ. Fluorescence intensity was used as a readout of relative cell count over time using the ‘plot profile’ function in ImageJ ^74^. Cell numbers were normalized to a baseline of “100 %” at the start of treatment (shown as time 0). Distribution of the bacterial population was quantified through use of the ‘radial profile’ ImageJ plugin. Radial distribution data were normalized by setting the field of view periphery as the baseline of “1-fold,” which we defined as 240 µm distance from the source. Images and videos shown were processed using the ‘enhance contrast’ function in ImageJ and adjusting intensity thresholds to normalize fluorescence intensity per cell across channels. For experiments with non-fluorescent cells, equivalent procedures were performed using phase contrast data and enumeration of cells over time using a Matlab-based tracking software ^6^.

#### Statistical Analyses

Data from replicate experiments were averaged and interpreted on the basis of their mean, standard error of the mean, and effect sizes. Effect sizes for data are indicated as Cohen’s *d* value:

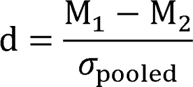

where:

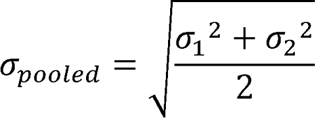

Where relevant, *p*-values were calculated using either one-sided or unpaired two-sided tests, with significance determined at *p*<0.05:

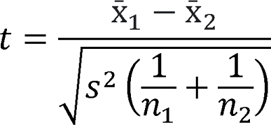

where:

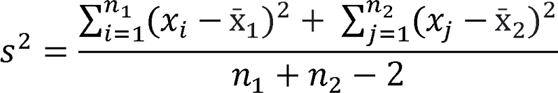

## Acknowledgements

Funding for this work was provided by NIAID under award numbers 1K99AI148587 and 4R00AI148587-03, and startup funding from Washington State University to AB. We thank Karen Guillemin (University of Oregon) for the Eppendorf Transjector used for the CIRA experiments. We thank Nikki Shariat (University of Georgia, Athens), Nkuchia Mikanatha and Pennsylvania NARMS and GenomeTrakr Programs, and Andreas Bäumler (University of California, Davis) for providing the *Salmonella* strains used in this work. Beamline 5.0.3 of the Advanced Light Source, a DOE Office of Science User Facility under Contract No. DE-AC02- 05CH11231, is supported in part by the ALS-ENABLE program funded by the National Institutes of Health, National Institute of General Medical Sciences, grant P30 GM124169-01.

All research on human and animal samples was performed in accordance with, and approval of, the Institutional Biosafety Committee and Institutional Animal Care and Use Committee at Washington State University.

## Author Contributions

A.B. and S.G. conducted the CIRA experiments. S.G. and Z.G. performed growth curve analyses with human serum and purified L-serine. A.B., S.G., and Z.G. performed the crystallographic analyses. T.A. performed the microgradient modeling. M.S. and M.J.H. performed the ITC experiments. All authors contributed to data analyses and writing of the manuscript.

## Movies

**Movie 1.** Representative CIRA experiments with Alexa Fluor 488 dye (left) and *S. enterica* Typhimurium IR715 treated with human serum (right). Video is shown at 1x speed and is also viewable at: https://www.youtube.com/watch?v=dyrQT2Ni5J8

**Movie 2.** Representative CIRA experiments comparing responses to human serum between *S. enterica* Typhimurium IR715 (*mplum*) and clinical isolates (*gfp*), as indicated. Videos depict responses over 5 minutes of treatment. Viewable at: https://youtu.be/dwtZtoisjrU

**Movie 3.** Representative CIRA experiments comparing responses to human serum between WT *S. enterica* Typhimurium IR715 (*mplum* in left and center panel) and chemotactic mutants *tsr* (*gfp* in center panel, *mplum* in right panel), and *cheY* (*gfp* in left and right panel). Videos depict responses over 5 minutes of treatment. Viewable at: https://youtu.be/O5zsEAqcJw8

**Movie 4**. Representative CIRA experiments comparing responses to 500 µM L-serine between *S. enterica* Typhimurium IR715 (*mplum*) and clinical isolates (*gfp*), as indicated. Videos depict responses over 5 minutes of treatment. Viewable at: https://www.youtube.com/watch?v=p0Tsp06ZHO8

**Movie 5.** CIRA experiments comparing response of *S.* Typhimurium IR715 to L-serine or to norepinephrine. Cells are unprimed or primed with NE, as indicated. Videos depict responses over 5 minutes of treatment. Viewable at: https://youtu.be/pUOlVjKYptc

**Movie 6.** Crystal structure of *S. enterica* Typhimurium Tsr LBD (PDB: 8FYV) and comparison with *E. coli* Tsr LBD crystal structure (PDB: 3ATP). Viewable at: https://youtu.be/OlowDhRLNhA

## Supplementary Information

### Supplemental Figures

**Fig. S1.**
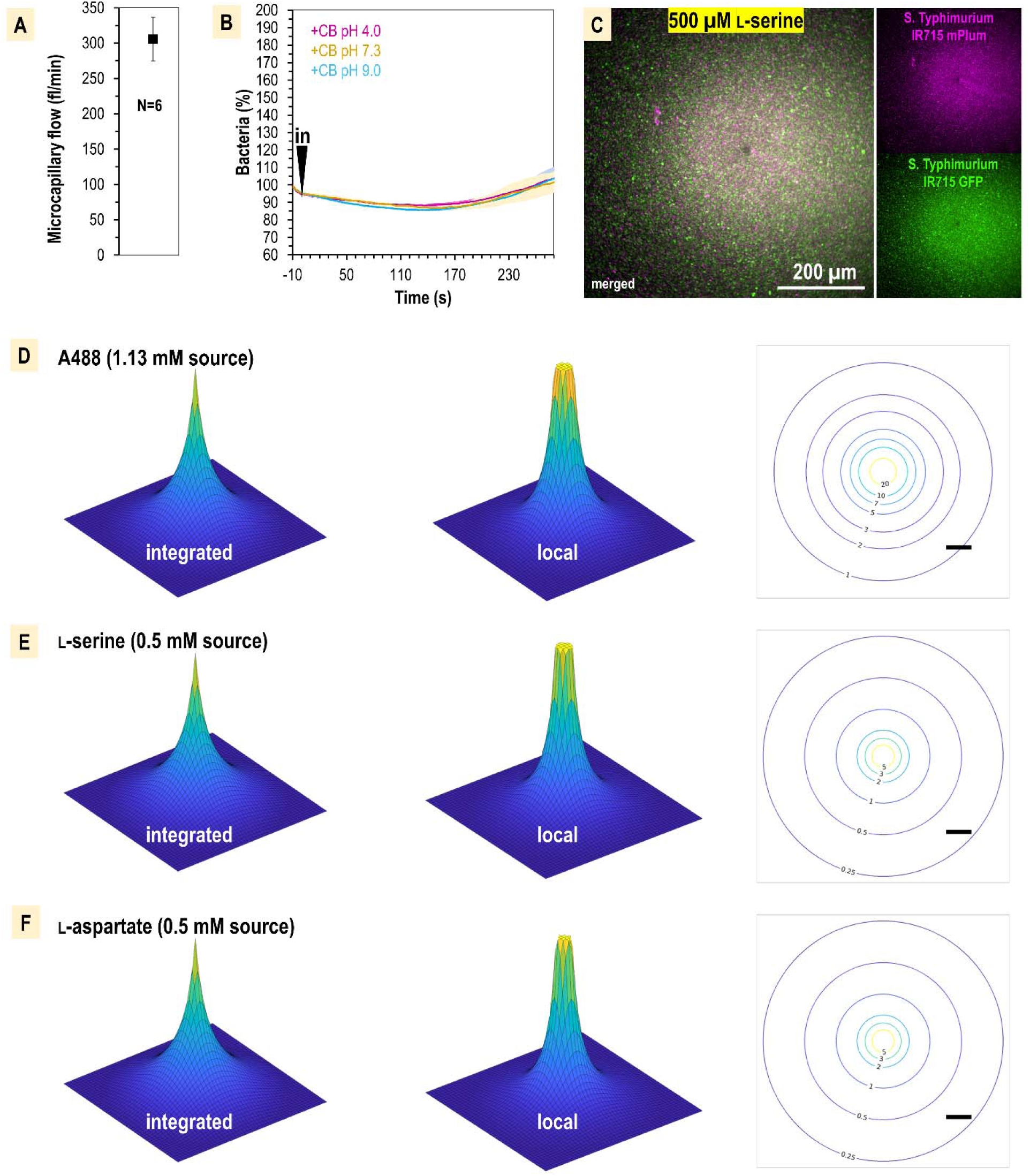
CIRA experimentation controls and microgradient modeling. A. Average injection flow of treatment solution from the glass microcapillary. Flow was measured through injection of known concentrations of methylene blue dye, diluted in CB, into a 50 µl pond of CB over 20 minutes. After treatments, the absorbance of solutions was measured at 665 nm and quantified based on a standard curve. Applying a compensation pressure of 35 hPa resulted in an average flow of 305.5 fl/min ± 31. B. CIRA localization in response to treatments of buffered CB of different pH. C. Comparison of responses of GFP versus mPlum strains using CIRA. D-F. Model of the CIRA microgradient for effectors relevant to this study. Topology maps of 1 mm x 1 mm size are shown for each microgradient in an ‘integrated’ format, which models what is seen by eye from the bottom-up view of the microscopy plane (left), or as the local concentration that would be experienced by a bacterium at a given distance (center). A plane through the center of the treatment sphere is shown (right), with relative concentrations experienced at a given distance expressed in nM units (scale bar is 100 µm).

**Fig. S2.**
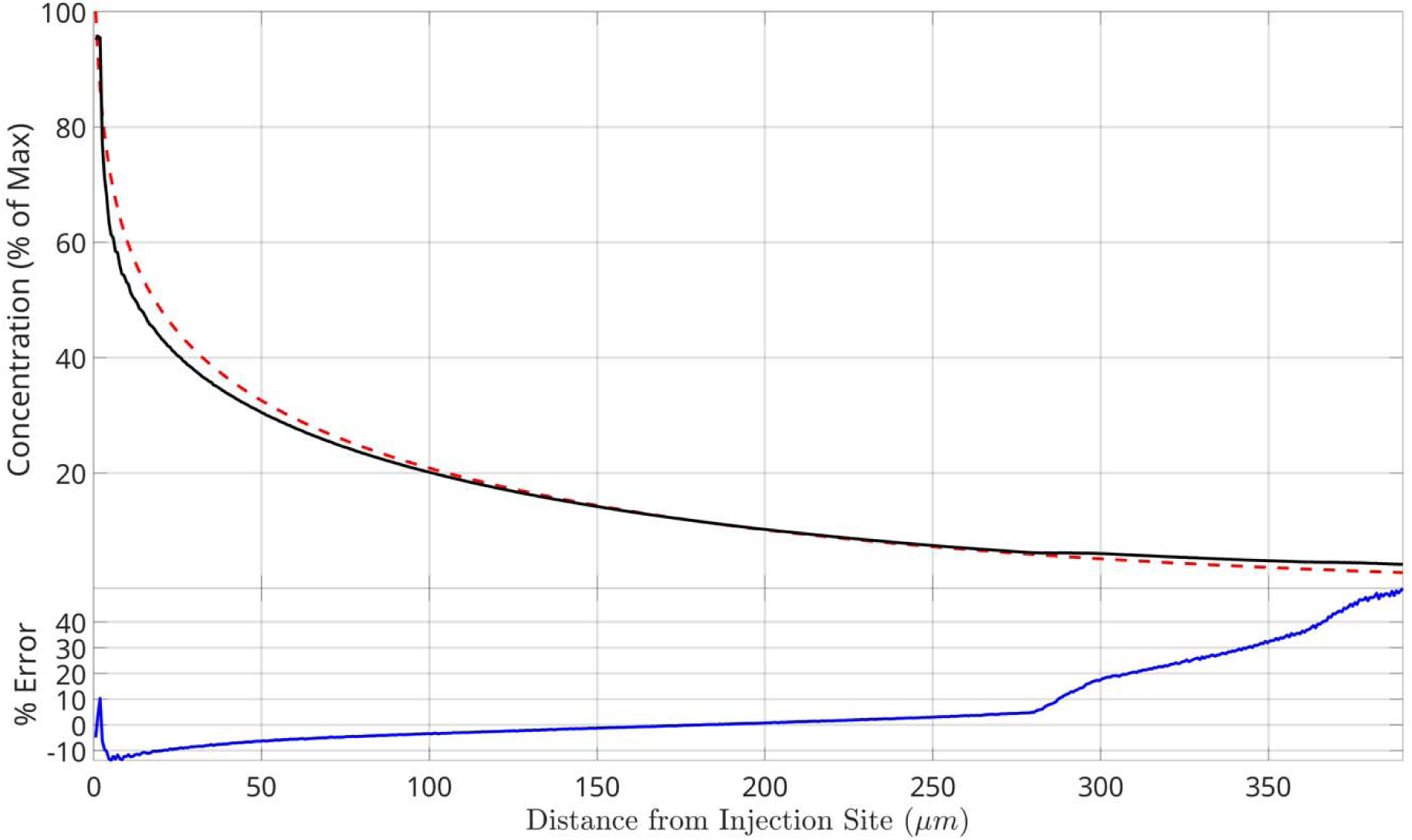
Comparison of CIRA diffusion modeling with experimental A488 data. Computed (red dashed) and measured (black solid) integrated concentrations of A488 as a function of distance from injection site for time t=120 sec. Both quantities are reported as a percentage of the concentration at the injection site (1.1 mM). The relative error (blue solid) is reported as the relative difference of the measured concentration to the computed concentration.

**Fig. S3.**
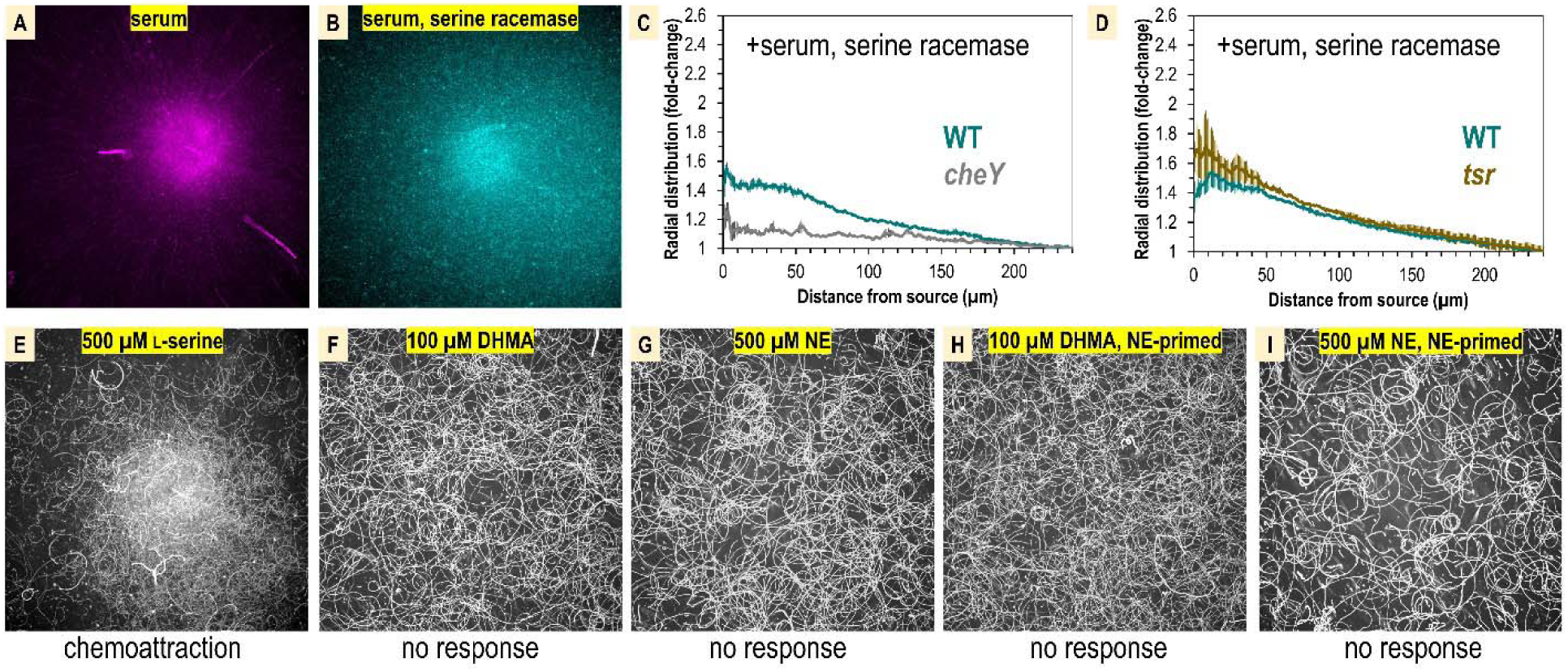
Comparison of *S.* Typhimurium IR715 chemotactic responses to human serum, human serum with serine racemase treatment, L-serine versus DHMA and NE. A-B. Shown are max projections from CIRA experiments over a 10 s time period following 300 s of treatments as shown. C-D. The radial distribution of bacteria in CIRA competition experiments with serine- racemase-treated human serum after 300 s is shown. E-I. Shown are max projections over 10 s following CIRA treatments, as indicated. The chemotactic response is noted below each experiment. Panels H-I utilized cells that were primed with 5 µM NE for 3 hours prior to experimentation.

**Fig. S4.**
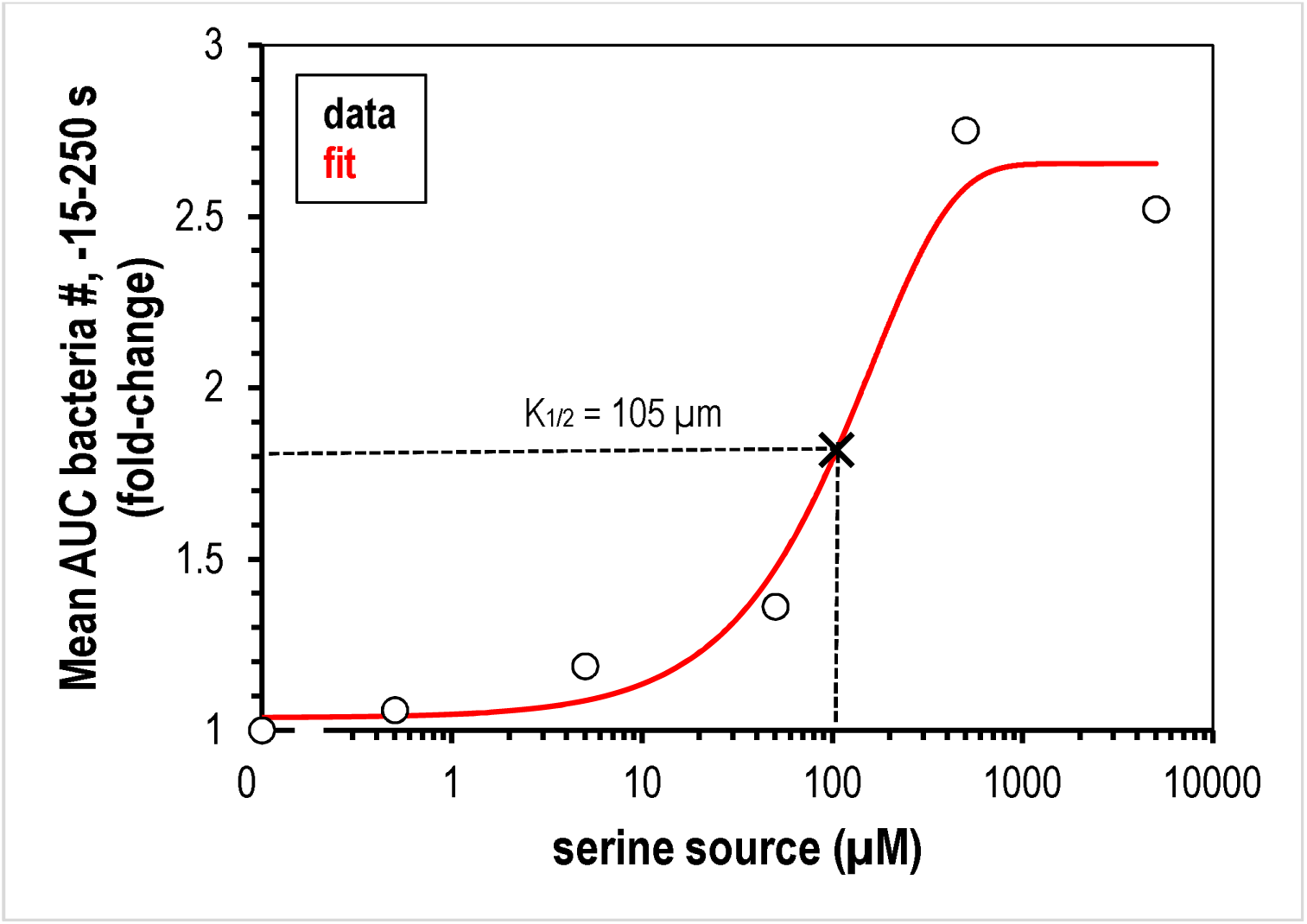
Calculation of L-serine source required for half-maximal chemoattraction response (K_1/2_). Data are mean AUC values of relative number of bacteria within 150 µm of the treatment location for the -15 s – 250 s range, represented as fold-change. Data are fit with an exponential curve (red line):

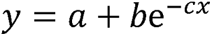 Where a=2.636193, b =-1.616338, and c=0.006282888. K_1/2_ is approximated to be 105 µm (x, dashed lines).

**Fig. S5.**
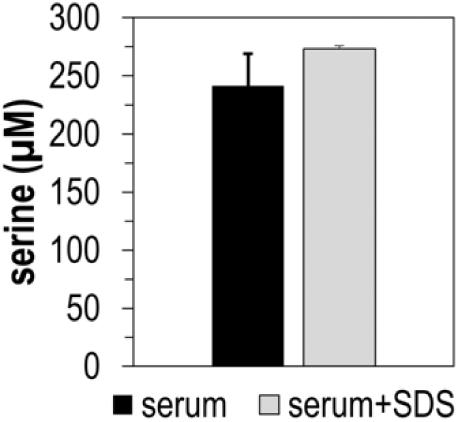
Total serine present in human serum samples, as determined by mass spectrometry. Treatment with 50 µg recombinant serine dehydratase (SDS) over 4 h did not decrease serine content in human serum.

**Fig. S6.**
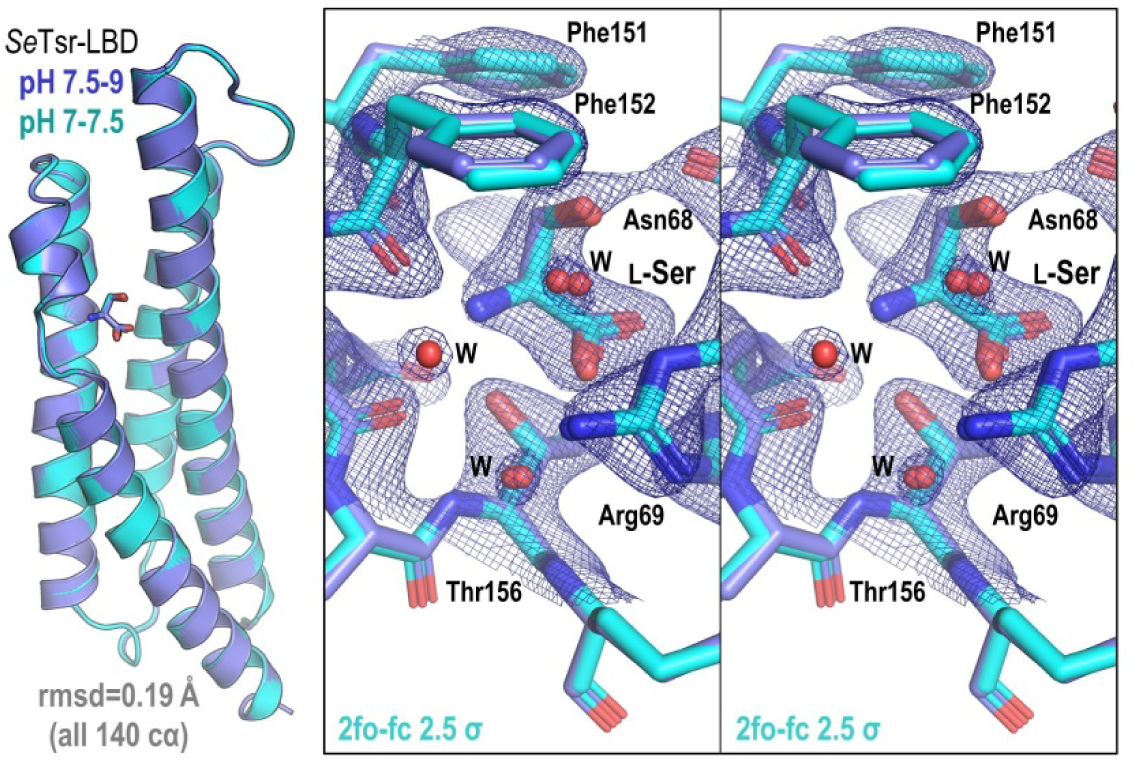
Comparison of *Se*Tsr LBD structures solved at different pH. Shown left is an overlay of *Se*Tsr at pH 7.5-9.7 (PDB: 8FYV, dark blue) and pH 7-7.5 (PDBL 8VL8, light blue). Shown right is a stereo image of the overlaid L-Ser binding site with electron density shown for the pH 7-7.5 structure as blue mesh.

## Supplementary Movies

**Movie S1.** CIRA experiments showing response of *S.* Typhimurium IR715 to DHMA. Cells are unprimed (left) or primed with NE (right). Viewable at: https://youtu.be/j4YL95QFCuI

**Movie S2.** Representative CIRA experiment showing response of *C. koseri* BAA-895 to human serum. Viewable at: https://youtu.be/9iMJz2OPbso

**Movie S3.** Representative CIRA experiment showing response of *E. coli* (pxS-gfp) MG1655 to human serum. Viewable at: https://www.youtube.com/watch?v=jq3cj9e52n4

## Supplementary Tables

**Table S1.**
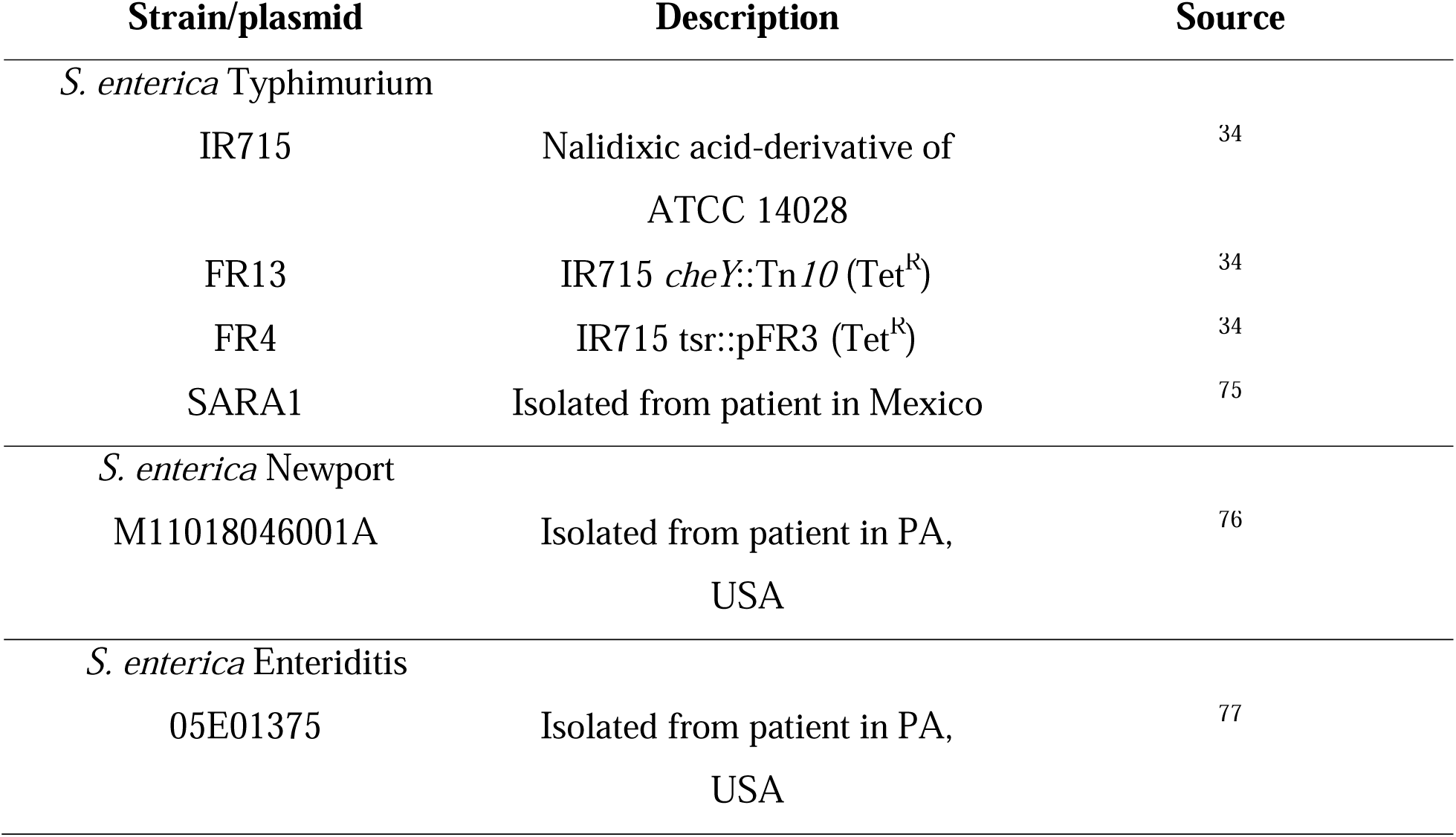
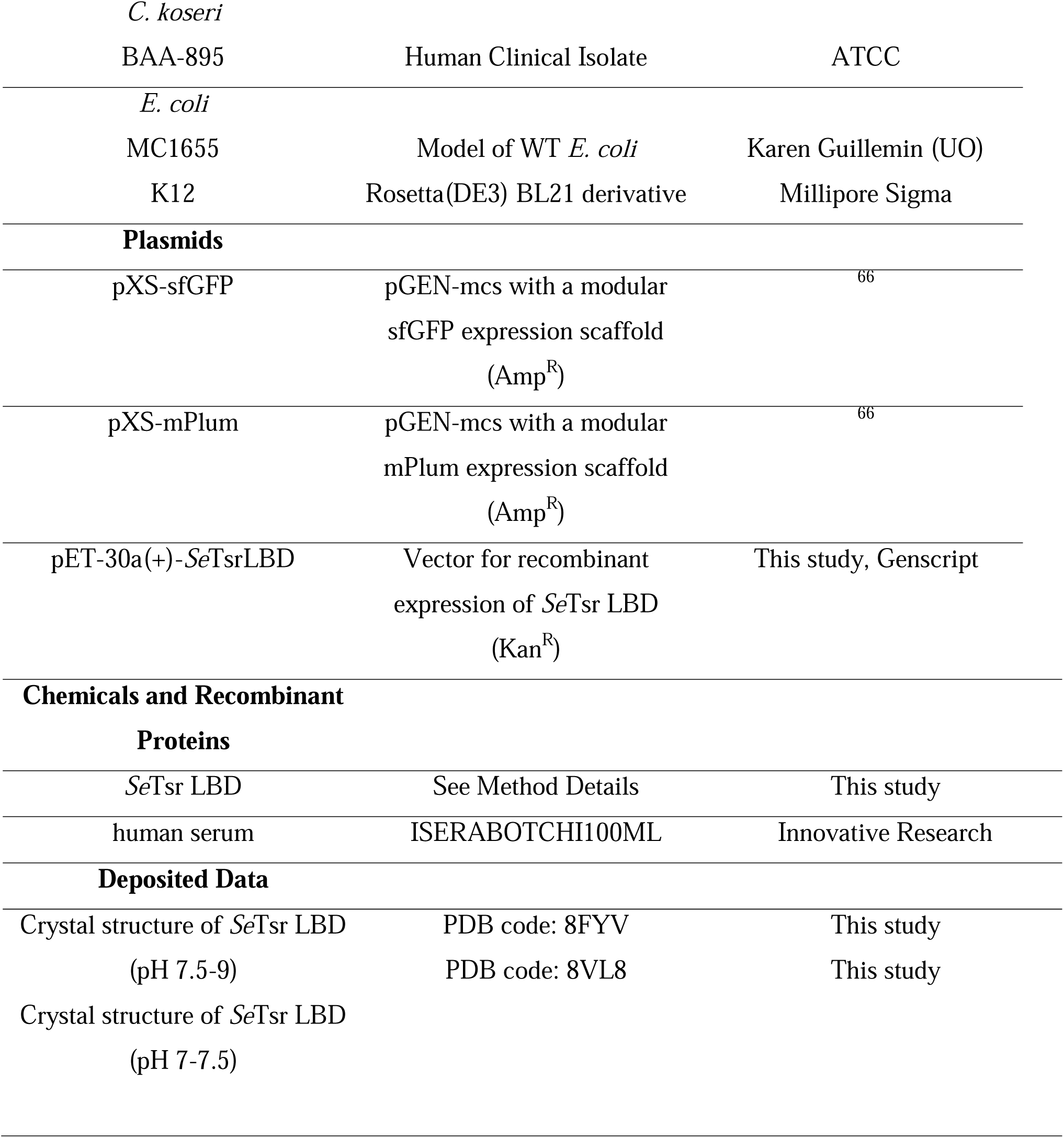
Bacterial strains and plasmids used in this study.

**Table S2.**
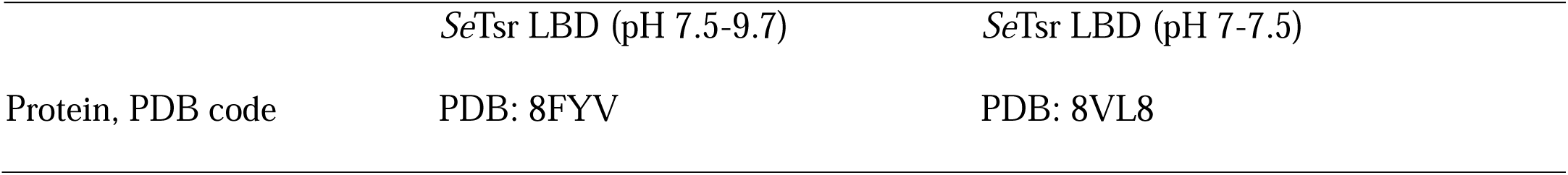
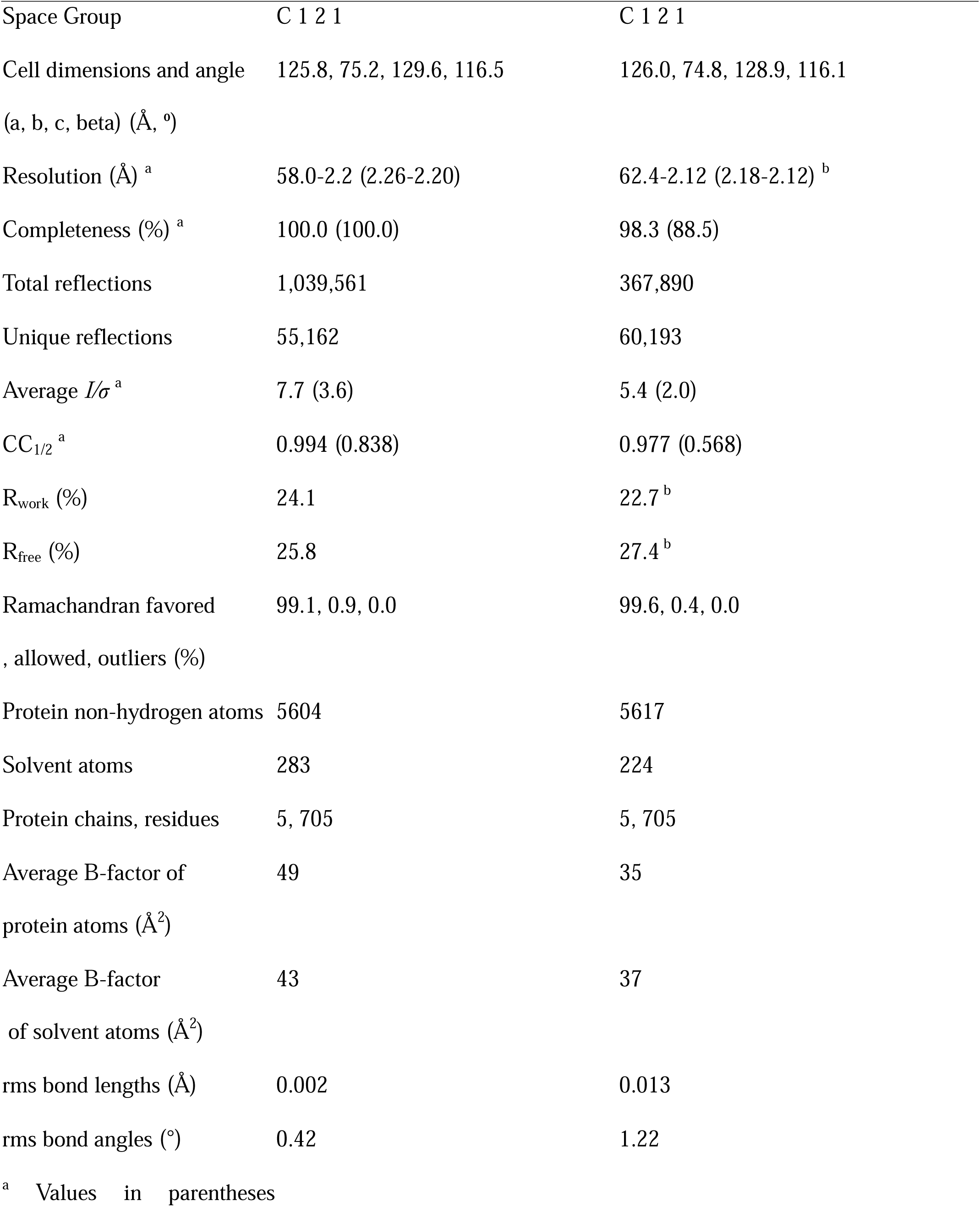
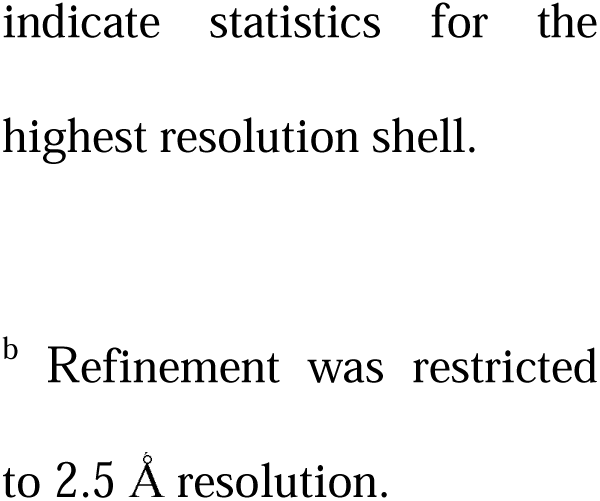
Summary of crystallographic statistics.

## Supplementary data

**Data S1.** Raw videos of CIRA experiments with DHMA and NE under various conditions. Experiment conditions are noted within ‘README.txt’ files. Data files can be downloaded at the following URL: http://tinyurl.com/CIRAdata1. If for some reason the data become inaccessible for download, they can be provided upon request to the correspond author.

**Data S2.** Database of Tsr homologue sequences.

